# Low-Magnitude Mechanical Signals Combined with Zoledronic Acid Reduce Musculoskeletal Weakness and Adiposity in Estrogen-Deprived Mice

**DOI:** 10.1101/2023.03.12.531571

**Authors:** Gabriel M. Pagnotti, Trupti Trivedi, Laura E. Wright, Sutha K. John, Sreemala Murthy, Ryan R. Pattyn, Monte S. Willis, Yun She, Sukanya Suresh, William R. Thompson, Clinton T. Rubin, Khalid S. Mohammad, Theresa A. Guise

## Abstract

Combination treatment of Low-Intensity Vibration (LIV) with zoledronic acid (ZA) was hypothesized to preserve bone mass and muscle strength while reducing adipose tissue accrual associated with complete estrogen (E_2_)-deprivation in young and skeletally mature mice. Complete E_2_-deprivation (surgical-ovariectomy (OVX) and daily injection of aromatase inhibitor (AI) letrozole) were performed on 8-week-old C57BL/6 female mice for 4 weeks following commencement of LIV administration or control (no LIV), for 28 weeks. Additionally, 16-week-old C57BL/6 female E_2_-deprived mice were administered ±LIV twice daily and supplemented with ±ZA (2.5 ng/kg/week). By week 28, lean tissue mass quantified by dual-energy X-ray absorptiometry was increased in younger OVX/AI+LIV(y) mice, with increased myofiber cross-sectional area of quadratus femorii. Grip strength was greater in OVX/AI+LIV(y) mice than OVX/AI(y) mice. Fat mass remained lower in OVX/AI+LIV(y) mice throughout the experiment compared with OVX/AI(y) mice. OVX/AI+LIV(y) mice exhibited increased glucose tolerance and reduced leptin and free fatty acids than OVX/AI(y) mice. Trabecular bone volume fraction and connectivity density increased in the vertebrae of OVX/AI+LIV(y) mice compared to OVX/AI(y) mice; however, this effect was attenuated in the older cohort of E_2_-deprived mice, specifically in OVX/AI+ZA mice, requiring combined LIV with ZA to increase trabecular bone volume and strength. Similar improvements in cortical bone thickness and cross-sectional area of the femoral mid-diaphysis were observed in OVX/AI+LIV+ZA mice, resulting in greater fracture resistance. Our findings demonstrate that the combination of mechanical signals in the form of LIV and anti-resorptive therapy via ZA improve vertebral trabecular bone and femoral cortical bone, increase lean mass, and reduce adiposity in mice undergoing complete E_2_-deprivation.

**One Sentence Summary:** Low-magnitude mechanical signals with zoledronic acid suppressed bone and muscle loss and adiposity in mice undergoing complete estrogen deprivation.

**Translational Relevance:** Postmenopausal patients with estrogen receptor-positive breast cancer treated with aromatase inhibitors to reduce tumor progression experience deleterious effects to bone and muscle subsequently develop muscle weakness, bone fragility, and adipose tissue accrual. Bisphosphonates (i.e., zoledronic acid) prescribed to inhibit osteoclast-mediated bone resorption are effective in preventing bone loss but may not address the non-skeletal effects of muscle weakness and fat accumulation that contribute to patient morbidity. Mechanical signals, typically delivered to the musculoskeletal system during exercise/physical activity, are integral for maintaining bone and muscle health; however, patients undergoing treatments for breast cancer often experience decreased physical activity which further accelerates musculoskeletal degeneration. Low-magnitude mechanical signals, in the form of low-intensity vibrations, generate dynamic loading forces similar to those derived from skeletal muscle contractility. As an adjuvant to existing treatment strategies, low-intensity vibrations may preserve or rescue diminished bone and muscle degraded by breast cancer treatment.

## Introduction

Breast cancer is among the most prevalent malignancies afflicting individuals in the United States [1, 2]. Approximately 80% of individuals with breast cancer have estrogen (E_2_) receptor–positive disease [3, 4], with the majority of cases occurring in early- to post-menopausal women. Untreated E_2_ receptor–positive breast cancer exhibits a high affinity to metastasize to bone, liver and lung [5, 6]. The skeleton is a favorable site of metastasis due to the high concentration and co-localization of growth factors; some of which are derived from the bone matrix, such as transforming growth factor-β (TGF-β) [7], and others are secreted by the tumor itself, including parathyroid hormone-related peptide [8, 9]. By virtue of proximity, these growth factors are locally available to drive tumor progression [7, 10–13]. Suppressing tumor advancement through pharmacologic and/or radiologic modalities is integral to prolonging patient lives [14–16]. Clinical strategies to inhibit breast cancer progression include selective E_2_-receptor (ER) modulators (e.g., tamoxifen) [17] for pre-menopausal patients and hormone-deprivation therapies (e.g., aromatase inhibitors; AI) [18] for postmenopausal patients. Third-generation AIs (e.g., letrozole) curtail peripheral estradiol synthesis from adrenal tissue by inhibiting aromatase production [19, 20]. These anti-E_2_ therapies are highly effective to decrease tumor-associated mortality and increase disease-free survival.

However, despite the advantages of suppressing E_2_ synthesis in postmenopausal patients treated with AIs, these drugs increase osteoclast-mediated resorption and subsequent bone destruction [21–23]. Such bone loss not only fuels tumor growth [24] but leads to increased release of TGF-β, which exacerbates muscle weakness [25–27] through dysregulation of ryanodine receptor-1 (RyR-1), a critical Ca^2+^-dependent Ca^2+^-release channel within the sarcoplasmic reticulum [28]. Denosumab, an antibody recognizing receptor-activator of nuclear-factor Κappa-β ligand (RANKL), is the first line of treatment prescribed to inhibit osteoclast-mediated bone resorption in postmenopausal women [29], as well as in patients with hormone receptor–positive breast cancer treated with AIs [30, 31]. Including bisphosphonates [e.g., zoledronic acid (ZA)] in the treatment regimen for breast cancer also effectively inhibits osteoclast activity and reduces cancer recurrence, improving survival in postmenopausal ER^+^ breast cancer patients [32]. Bisphosphonates prevent adhesion of metastatic cells to bone and induce osteoclast apoptosis *in vivo* [33] and *in vitro* [34, 35]. Thus, clinical challenges persist following treatment with AIs, including muscle weakness [36], progressive adiposity, and imbalanced bone remodeling [37, 38]. As these toxicities reduce quality of life [39], lead to poor patient adherence, and result in treatment discontinuation [40–42], novel approaches are needed to reduce the musculoskeletal burden on these individuals to ultimately maintain adherence to AI-mediated therapies, and to increase survival.

Daily physical activity is universally recommended as an alternative or supplemental therapy in standards-of-care to prevent bone and muscle loss [43–45], including in clinical cancer treatment guidelines. Exercise is a known to have anabolic effects on bone and muscle while suppressing fat formation [46, 47]. Additionally, exercise reduces inflammation [48] and regulates glucose metabolism. Despite its well-recognized benefits and patient willingness, breast cancer patients often find it difficult to participate in typical exercise regiments due to decreased energy and muscle strength [49]. Cancer patients also experience heightened pain and frailty, thus incorporating strenuous loads, such as those induced by exercise, may elevate fracture risk, therein precipitating the fractures that the exercise was intended to prevent [50, 51].

As an alternative to the high-intensity loads generated by daily locomotion let alone physical activity or exercise regimens [52], low-intensity vibration (LIV) provides a low magnitude (<1 g acceleration, where 1 g = Earth’s gravitational field or 9.8 m/s^2^) mechanical signal delivered through vertical (uniaxial) acceleration at a physiologically high frequency (∼20-100 Hz) but low-magnitude force that is anabolic to bone in both translational and clinical studies [53, 54]. LIV does not require strenuous engagement by the patient, providing a fraction of the dynamic mechanical energy used in postural muscle contraction [55] to maintain upright balance, which is itself a significant contributor to bone homeostasis and wanes with age [55, 56] or as a result of dystrophy or disease [54]. We previously showed that LIV safely preserves bone and reduces tumor burden in murine ovarian and myeloma cancer models [57, 58]. Additionally, LIV inhibits adiposity in mice fed a high-fat diet [59, 60] and increases the density of muscle-bound satellite cells in ovariectomized mice [61]. In mesenchymal stem cells, LIV induces osteogenic differentiation while attenuating adipogenesis [62, 63]. We’ve also shown that the timing of LIV dosing affects outcomes where delivering multiple bouts of LIV separated by a rest period results in beneficial effects in both bone tissue [59] and cells [64]. We recently showed that direct application of LIV to MDA-MB-231 breast cancer cells resulted in decreased invasion and impaired the ability to secrete paracrine factors that promote osteoclast differentiation [65]. In the clinic, LIV improved bone formation in young women with low bone mineral density [66] and in adolescent patients with anorexia nervosa [67] and cerebral palsy [68], as well as increased t-scores of childhood cancer survivors after 1 year of LIV treatment when compared with placebo-treated patients [69].

As LIV has protective effects on bone and muscle observed in preclinical studies of high-fat diet and cancer models and in patients with cancer remission and suppressed E_2_ production, we hypothesized that daily administration of LIV would preserve critical bone and muscle endpoints in the context of treatment-induced E_2_-deprivation. In the current study, we used mouse models designed to simulate the extreme bone loss, muscle weakness, and increased adiposity experienced by patients undergoing AI treatment to induce complete E_2_ deprivation.

## Results

### Experiment I: Determine the Effects of Low Intensity Vibration in Young Estrogen Deprived Mice

#### LIV Increased Trabecular Bone Microarchitecture and Histomorphometric Markers

Trabecular bone volume fraction (Tb.BV/TV) and trabecular connectivity density (Tb.Conn.D) in L5 vertebrae of OVX/AI+LIV(y) mice were greater (p < 0.047 and p < 0.017, respectively) than in OVX/AI(y) mice (Fig.1A). Although Tb.BV/TV and Tb.Conn.D were greater in L4 vertebrae of OVX/AI+LIV(y) mice than in OVX/AI(y) mice, these values were not significant. Respective increases in trabecular thickness, trabecular number, and trabecular bone mineral density (Tb.Th, Tb.N, and Tb.BMD, respectively) were observed in the L5 and L4 of OVX/AI+LIV(y) mice relative to OVX/AI(y) mice as well, but these values were not significant. (Note: Data from microCT analyses are presented in their entirety in Supplementary Table 2. The number of osteoclasts (N.OC/BS) and osteoclast surface area (OC.S/BS) in OVX/AI+LIV(y) L5 vertebrae were lower (p < 0.02 and p < 0.002, respectively) than in OVX/AI mice (Fig.1B). Dynamic histomorphometric analysis of vertebral trabecular bone indicated that bone formation rate (BFR/BS) was greater (p < 0.01) in OVX/AI+LIV(y) mice relative to OVX/AI(y) mice (Fig.1B). The number of osteoblasts (OB/BS) in OVX/AI+LIV(y) mice was greater (p < 0.01) than in OVX/AI(y) mice (Fig.1B). Dynamic histomorphometry of femoral trabecular bone showed an increase (p < 0.001) in bone formation rate (BFR/BS) and in mineral apposition rate (MAR) (p < 0.003) of OVX/AI+LIV(y) mice relative to that of OVX/AI(y) mice (Fig.1C). No differences were observed in N.OC/BS or OC.S/BS across the femoral metaphysis.

**Figure 1:**
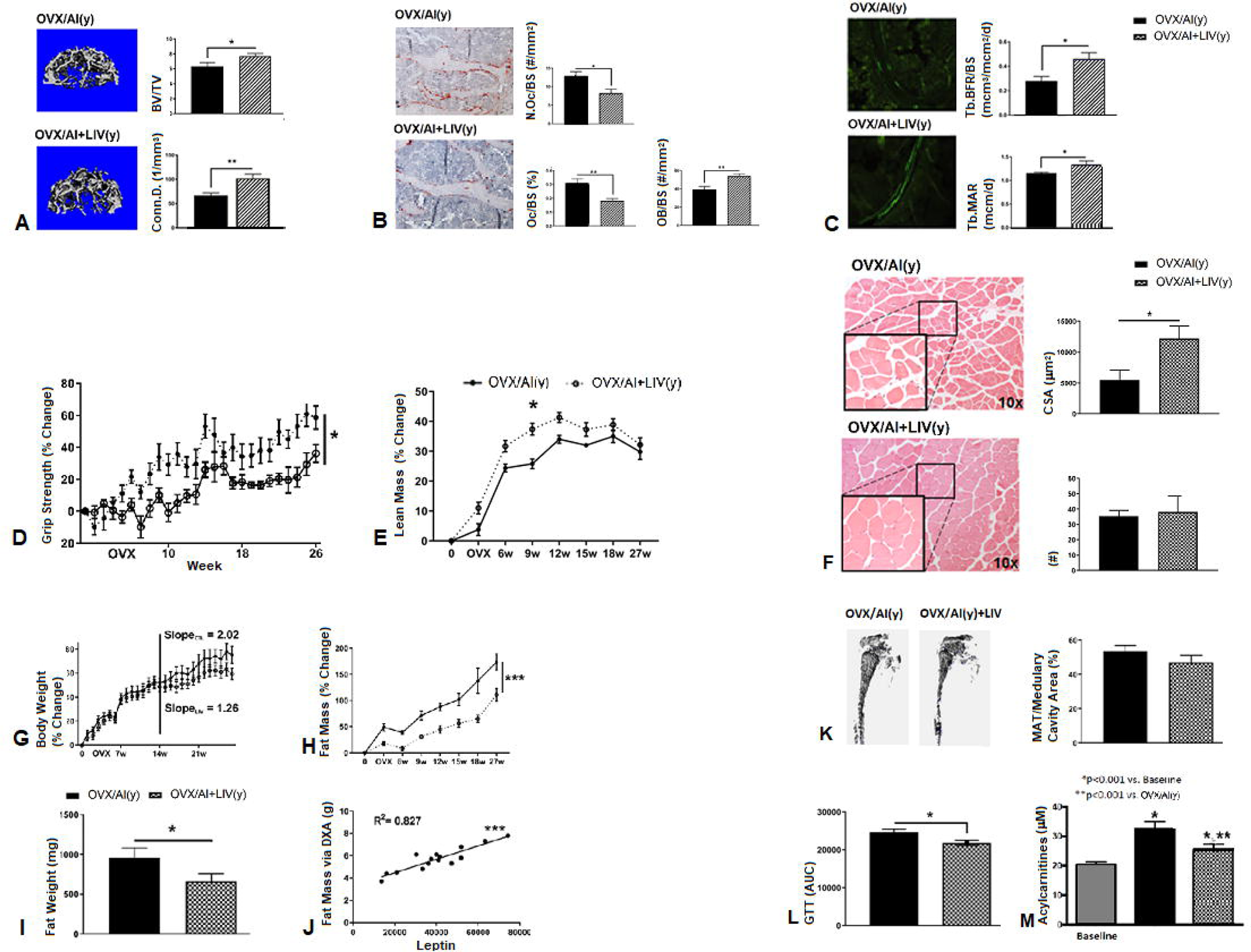
Experiment I: LIV Improves Bone and Muscle While Reducing Adiposity in Young, E_2_-Deprived Mice. Results from experiment I (the younger mouse cohort) are shown for differences in (**A.**) trabecular bone microarchitecture and (**B**.) static and (**C**.) dynamic histomorphometry. Significant decreases in N.OC/BS and OC.S/BS were observed in OVX/AI+LIV(y) mice as confirmed by tartrate-resistant acid phosphatase–stained sections exhibiting a reduced number of positively stained osteoclasts. Histomorphometry of OVX/AI+LIV(y) L5 vertebral body showed significant increases in trabecular BFR/BS and OB/BS, as indicated in the images (magnification 10×) by the greater fluorescent double-labeled surface area, as compared to OVX/AI(y). Differences between OVX/AI(y) and OVX/AI+LIV(y) are shown in terms of **(D**.) muscle function (i.e. forelimb grip strength) and (**E**.) % whole body lean mass (as quantified by DXA) to accompany analysis of (**F**.) myofiber cross-sectional area and number. (**G**.) Whole-body weight was tracked weekly with the rate of increase performed from week 14 until euthanasia. Adiposity was quantified by (**H**.) whole-body fat mass (as quantified by DXA) and (**I**.) perigonadal fat pad weight. (**J**.) % fat mass was significantly correlated to serum leptin. (**K**.) Osmium tetroxide staining was used to quantify marrow adiposity. Metabolism was quantified by (**L**.) glucose tolerance testing and analyzing area under the curve, while (**M**.) metabolomics was performed on serum to quantify multiple-sized free-fatty acid chains. Abbreviations: OVX, ovariectomy; AI, aromatase inhibitor; LIV, low-intensity vibration; [OVX/AI(y)]. *p < 0.05; **p < 0.01; ***p < 0.001; ****p < 0.0001.

#### LIV Increased Muscle Strength and Function Following Estrogen Deprivation

From 8 weeks of age (after OVX) until euthanasia (32-week-old), longitudinal change in forelimb grip strength measurements relative to baseline readings were significantly greater (p < 0.05) in OVX/AI+LIV(y) mice than in OVX/AI(y) mice (Fig.1D). By 26 weeks after OVX, OVX/AI+LIV(y) mice exhibited greater (p < 0.03) grip strength than OVX/AI(y) mice (Fig.1D). Throughout the study, significant effects were observed as a result of treatment with LIV (p < 0.05), time (p < 0.0001), and between subjects (p < 0.0001), and an interaction effect (p < 0.0001) was also observed. Tetanic stimulation of the EDL at euthanasia exhibited an interaction (p < 0.01), time (p < 0.001), and subject-related (p < 0.001) effect on muscle fatigue between OVX/AI(y) and OVX/AI+LIV(y) mice. Although there were time and subject-related effects, contractility of the EDL did not exhibit any differences in muscle specific force between OVX/AI(y) and OVX/AI+LIV(y) mice. Longitudinal DXA scans showed that OVX/AI+LIV(y) mice had greater (p < 0.03) lean mass relative to OVX/AI(y) mice by week-28 (Fig.1E). Muscle from LIV treated mice had increased myofiber size with decreased spacing between individual fibers (Fig.1F). Histologic sections of OVX/AI+LIV(y) quadratus femora showed increased myofiber cross-sectional (CSA; p < 0.034) by 24 weeks compared with OVX/AI(y) mice (Fig.1F). No differences in the number of myofibers were observed between the two groups (Fig.1F).

#### LIV Suppressed Body Weight Gains, Adiposity, and Increased Glucose Metabolism Associated with Estrogen Deprivation

Longitudinal measurements of body weight in young mice in experiment I showed no differences between the OVX/AI(y) and OVX/AI+LIV(y) groups (Fig.1G). However, linear regression analysis from week-14 to week-28 after OVX showed a significant difference between slopes, with OVX/AI+LIV(y) mice gaining weight at half the rate of OVX/AI(y) mice (Fig.1G). OVX/AI+LIV(y) mice had lower (p < 0.0001) fat mass relative to OVX/AI(y) mice by week-28 (Fig.1H). The weight of fat pad harvested from the perigonadal depots at euthanasia was significantly (p < 0.05) lower in OVX/AI+LIV(y) mice than in OVX/AI(y) mice after 28 weeks of treatment with LIV (Fig.1I). Serum leptin exhibited a decreasing trend in OVX/AI+LIV(y) mice at 24 weeks after OVX that, when compared with total fat mass at euthanasia, resulted in a significant correlation (R^2^ = 0.827, p < 0.001; Fig.1J). Marrow adipose tissue quantified from the tibiae of younger mice was 6.5% lower in OVX/AI+LIV(y) mice than in OVX/AI(y) mice when normalized to medullary cavity volume, but this was not significantly different (Fig.1K). Blood glucose readings in young mice were recorded using a glucometer at 0, 15, 30, 60, and 120 minutes from blood draw at the tail vein. OVX/AI+LIV(y) mice exhibited greater serum blood glucose tolerance (p = 0.05) than OVX/AI(y) mice, as quantified by the area under the curve (Fig.1L). Analysis of serum fatty acid oxidation intermediates (acylcarnitines) in young mice prior to and 28 weeks after commencing E_2_ deprivation identified significant increases in short-chain and medium-chain acylcarnitines. Specifically, 28 weeks of E_2_ deprivation induced greater (p < 0.05) circulating short-chain acylcarnitine (consisting of 2-7 carbon chains) concentrations compared with baseline (Fig.1M). Moreover, OVX/AI+LIV(y) mice at 28 weeks had significant attenuation (p < 0.001) of the C2-C7 acylcarnitine, and this did not return to pre–E_2_ deprivation levels. Among the C2-C7 acylcarnitine groups, acetylcarnitine (C2) and hexanoylcarnitine (C6) were the significantly changed species contributing to the changes observed at 28 weeks of E_2_ deprivation in both groups in experiment I (Supplementary Fig.1). Medium-chain (C8-C14) acylcarnitines significantly increased (p < 0.001) after 28 weeks of E_2_ deprivation, and these levels did not decrease in OVX/AI+LIV(y) mice relative to OVX/AI(y) mice (Supplementary Fig.2). No differences in long-chain (C16-C22) species were observed between groups (Supplementary Fig.2).

### Experiment II: Determine the Effects of Low Intensity Vibration in Combination with Zoledronic Acid in Skeletally Mature Estrogen Deprived Mice

#### Bone Mineral Density and Cortical and Trabecular Microarchitecture are Enhanced When LIV is Administered in Combination with ZA

(Note: MicroCT data are presented in their entirety in Supplementary Table S2). DXA-obtained whole-body BMD in the older mice subject to E_2_ deprivation (OVX/AI) was decreased (p < 0.0001) compared with SH mice (Fig.2A). OVX/AI+LIV mice exhibited no differences relative to OVX/AI mice. OVX/AI+ZA mice exhibited increased whole-body BMD (p < 0.0001) relative to OVX/AI mice but lower whole-body BMD (p < 0.01) than SH mice (Fig.2B). However, whole-body BMD was increased (p < 0.0001) in OVX/AI+LIV+ZA mice compared with OVX/AI mice, an increase when compared with SH mice (Fig.2A). BMD analysis of the lumbar spine in the skeletally mature mice using DXA exhibited time, group, and interaction effects (p < 0.0001; Fig.2B). OVX/AI and OVX/AI+LIV mice exhibited a reduction (p < 0.0001) in lumbar BMD relative to SH mice as early as 4 weeks after OVX (Fig.2B) and this effect continued throughout the study. However, at 4 weeks after OVX, lumbar BMD in OVX/AI+ZA mice was lower (p < 0.01) than in SH mice. Lumbar BMD values in OVX/AI+ZA mice were increased compared with OVX/AI and OVX/AI+LIV mice (Fig.2B). Additionally, OVX/AI+LIV+ZA mice exhibited no differences in lumbar BMD 4 weeks after OVX relative to SH mice, which was greater (p < 0.0001) than the difference observed between OVX/AI and OVX/AI+LIV mice (Fig.2B). By 15 weeks after OVX, BMD of the lumbar vertebrae in OVX-AI+ZA mice was higher (p < 0.001) than in OVX/AI mice but only mildly greater (p < 0.01) than in OVX/AI+LIV mice (Fig.2B). However, lumbar BMD of OVX/AI+LIV+ZA mice was greater (p < 0.0001) than in OVX/AI mice at 15 weeks after OVX (Fig.2B).

**Figure 2:**
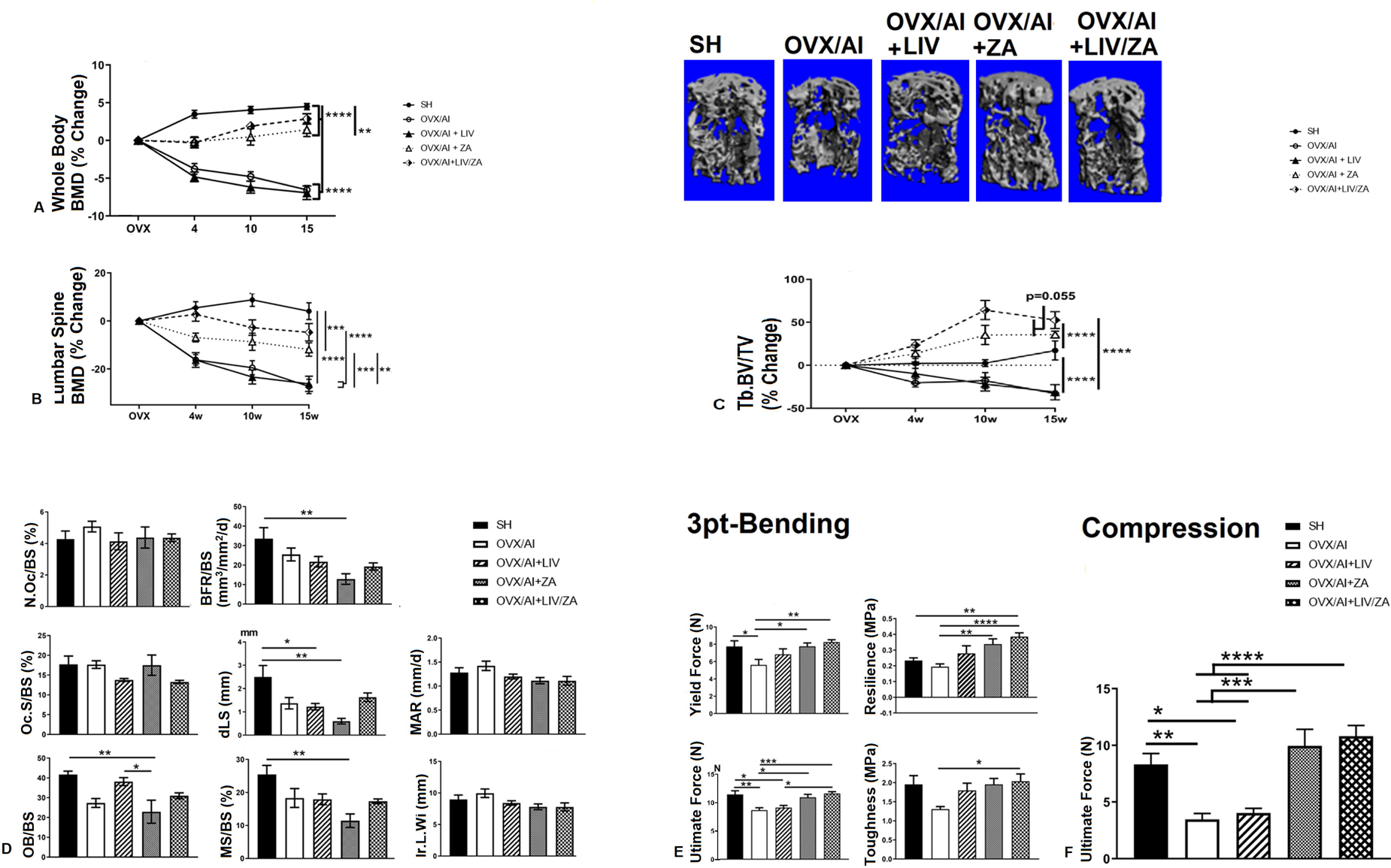
Bone mineral density (BMD), microarchitecture, mechanical, and structural properties of mice in experiment II (older cohort). Longitudinal dual-energy x-ray absorptiometry (DXA) scans were performed to quantify (A.) whole-body and (B.) lumbar spine BMD for regional measurements. Significant loss of whole-body and lumbar BMD was observed in untreated and LIV-treated OVX/AI mice. Whole-body BMD was preserved in OVX/AI+ZA mice, and this effect was improved even further when LIV was included in the treatment regimen (i.e. OVX/AI+LIV+ZA), restoring BMD near to that of SH mice. (C.) Longitudinal micro-computed tomography scans of trabecular-rich L5 vertebral body showed significantly decreased BV/TV in OVX/AI-mice. OVX/AI+ZA mice maintained positive accrual of bone throughout the study, exceeding values measured from SH mice. These findings improved further with the addition of LIV to the treatment regimen (i.e. OVX/AI+LIV+ZA). (D.) Static histomorphometry did not demonstrate any differences in N.OC/BS or OC.S/BS; however, OVX/AI+LIV and OVX/AI+LIV+ZA exhibited a trend of reduced OC.S/BS in contrast to OVX/AI vehicle controls. OB/BS was decreased in the vertebrae of both OVX/AI and OVX/AI+ZA mice, and mice treated with LIV either with or without ZA had preserved trabecular bone-bound OB. Dynamic histomorphometry exhibited significant loss of BFR/BS, dLS, and MS/BS in OVX/AI+ZA mice compared with SH mice; however, combining LIV with ZA (i.e., in OVX/AI+LIV+ZA mice) induced a measure of protection. (E.) Micro-CT data highlighted that treatment with ZA improved BV/TV, and treatment with LIV further increased BV/TV, as visualized in 3-dimensional reconstructions. (F.) Functional testing of the L5 vertebrae via compression testing resulted in significantly reduced ultimate forces necessary to bring the bones to failure in OVX/AI mice without treatment; ZA significantly improved these outcomes. Near identical outcomes were observed when femora were subjected to 3-point bending, where untreated OVX/AI mice exhibited significantly reduced material and tissue level properties that increased after treatment with ZA and further increased with the addition of LIV. Abbreviations: OVX, ovariectomy; AI, aromatase inhibitor; LIV, low-intensity vibration; SH, sham-OVX and saline vehicle (control); ZA, zoledronic acid; w, weeks after OVX; Tb, trabecular; BFR/BS, bone formation rate; OB/BS, number of osteoblasts; N.OC/BS, number of osteoclasts; OC.S/BS, osteoclast surface area; dLS, double-label fluorescent surface area; MS/BS, mineralizing surface area; Ir.L.Wi, Interlabel Width; MAR, Mineral Apposition Rate; BV/TV, bone volume fraction. *p < 0.05; **p < 0.01; ***p < 0.001; ****p < 0.0001.

L5 and L4 vertebrae in OVX/AI mice exhibited a significant (p < 0.0001) decrease in Tb.BV/TV compared with those of SH mice by 15 weeks after OVX (Fig.2C). Tb.BV/TV was decreased (p < 0.0001) in L5 and L4 of OVX/AI mice relative to SH mice but was increased (p < 0.0001) in L5 and L4 of OVX/AI+ZA mice (Fig.2C). In OVX/AI+LIV+ZA mice, Tb.BV/TV increased (p < 0.0001) compared with untreated OVX/AI mice (Fig.2C). Interaction (p < 0.0001) and time (p < 0.003) effects were found between groups throughout the study as a function of percentage change in L5 Tb.BV/TV (Fig.2C). Untreated OVX/AI mice had decreased Tb.BV/TV (p < 0.0001) relative to SH mice (Fig.2C). OVX/AI+ZA mice exhibited an increase (p < 0.0001) in Tb.BV/TV relative to OVX/AI mice, and in OVX/AI+LIV mice, Tb.BV/TV was greater (p < 0.0001) than in untreated OVX/AI mice; a similar effect was also observed (p < 0.01) in OVX/AI+LIV+ZA mice (Fig.2C). These findings were paralleled by similar differences in Tb.N and Tb.Th, each showing decreases (p < 0.05 and p < 0.01, respectively) in OVX/AI mice relative to SH mice and increases (p < 0.05 and p < 0.001, respectively) in mice treated with ZA. OVX/AI+LIV+ZA mice exhibited increased Tb.N and Tb.Th (p < 0.001 and p < 0.0001, respectively) compared with OVX/AI mice, as well as increases relative to OVX/AI+ZA mice (p < 0.05 and p < 0.05, respectively).

Femoral Ct.BA/TA exhibited significant interaction, time, and treatment (ZA and LIV)-related effects (p < 0.0001) in older mice. Ct.BA/TA decreased (p < 0.0001) in OVX/AI mice relative to SH mice by 15 weeks after OVX. Although OVX/AI+LIV mice did not show improved Ct.BA/TA, Ct.BA/TA was increased (p < 0.001) in OVX/AI+ZA mice (Table S2). OVX/AI+ZA mice had increased (p < 0.0001) Ct.BA/TA, and OVX/AI+LIV mice had increased (p < 0.0001) Ct.BA/TA as well. Ct.BA/TA in OVX/AI+LIV+ZA mice increased (p < 0.0001) compared with that of OVX/AI mice, thereby improving on treatment with ZA alone. Corresponding decreases in Ct.Th (p < 0.0001) and Ct.pMOI (p < 0.05) were observed in OVX/AI mice compared with SH mice (Table S2). OVX/AI+ZA mice showed increased (p < 0.001) Ct.Th relative to OVX/AI mice, and Ct.Th increased even further (p < 0.0001) in OVX/AI+LIV+ZA mice. Ct.pMOI was not statistically different in OVX/AI+ZA or OVX/AI+LIV+ZA mice relative to OVX/AI or SH mice.

#### Decreased Bone Formation and Mineralization in Estrogen Deprived Mice Due to ZA is Attenuated in LIV-Treated Mice

BFR/BS was lower (p < 0.01) in OVX/AI+ZA mice than in SH mice, and OVX/AI+LIV+ZA mice had lower BFR/BS than SH mice (p < 0.06; Fig.2D). Double-label fluorescent surface area (dLS) of untreated OVX/AI mice was decreased compared to SH mice, but this was not significant (p = 0.08) (Fig.2D). OVX/AI+LIV and OVX/AI+ZA mice each showed reduced (p < 0.03 and p < 0.002, respectively) dLS compared with SH mice (Fig.2D); however, dLS of OVX/AI+LIV+ZA mice, although not statistically different from SH mice, was 2 times greater than that observed in OVX/AI+ZA mice (Fig.2D). Mineralizing surface area (MS/BS) was significantly reduced (p < 0.002) in OVX/AI+ZA mice relative to SH mice, but to a lesser degree (p = 0.07) in OVX/AI+LIV+ZA mice (Fig.2D). OB/BS was significantly decreased (p < 0.01) in OVX/AI+ZA mice compared with SH mice and was also lower (p < 0.03) compared with OVX/AI+LIV mice (Fig.2D). No differences were observed in OC.S/BS or N.OC/BS. Von Kossa-McNeil–stained sections of the femoral metaphysis showed a significant reduction (p < 0.05) in osteoblast number in OVX/AI+ZA mice relative to untreated OVX/AI mice but no differences were observed in OVX/AI+LIV+ZA mice. Sections stained with tartrate-resistant acid phosphatase exhibited no differences in the number of osteoclasts or osteoclast surface area between groups; however, there were fewer osteoclasts in OVX/AI+LIV mice at a similar proportion to that observed with OVX/AI+ZA mice relative to untreated OVX/AI mice.

#### The Mechanical Properties of Vertebral and Diaphyseal Bone Degraded by Estrogen Deprivation Are Normalized with ZA but Improved Further When Combined with LIV

(Note: Mechanical Testing of Bone: Mechanical testing data are presented in their entirety in Supplementary Table 3). Femora of OVX/AI mice exhibited a reduction (p < 0.001) in ultimate force compared to SH mice, but ultimate force was increased (p < 0.001) in OVX/AI+ZA mice relative to OVX/AI mice. Ultimate force in OVX/AI+LIV+ZA mice improved (p < 0.012) relative to OVX/AI mice (Fig.2E). Similarly, stiffness was also reduced (p < 0.002) in OVX/AI mice relative to SH mice, but no significant differences were observed in OVX/AI+LIV, OVX/AI+ZA, or OVX/AI+LIV+ZA mice relative to either SH or OVX/AI mice (Fig.2E). Tissue-level properties indicated similar decreases in yield stress, ultimate stress, resilience, and toughness in OVX/AI mice compared with SH mice. Improvements (p < 0.007) were observed in OVX/AI+ZA mice compared with OVX/AI mice. Nonsignificant increases were observed in resilience (47%), toughness (38%) (Fig.2E), and yield stress (18%) in OVX/AI+LIV mice relative to untreated-OVX/AI mice. OVX/AI+LIV+ZA mice exhibited increased resilience (p < 0.0001), toughness (p < 0.04) (Fig.2E), yield stress (p < 0.04), and ultimate stress (p < 0.02) compared with OVX/AI mice (Table S3).

L5 vertebrae of OVX/AI mice exhibited a decrease (p < 0.002) in ultimate force compared with that of SH mice. Ultimate force was increased in L5 (p < 0.0001) and L4 (p < 0.01) vertebrae of OVX/AI+ZA mice compared with untreated OVX/AI mice (Fig.2F). Ultimate force of L5 in OVX/AI+LIV+ZA mice improved (p < 0.0001) relative to OVX/AI mice, a 43% increase compared with OVX/AI+ZA (Fig.2F). L4 vertebrae from OVX/AI+LIV+ZA mice showed an improvement (p < 0.0001) compared with OVX/AI mice, which was greater (p < 0.02) than in E_2_-replete SH mice (Table S3). L4 and L5 vertebrae exhibited no differences in length between groups.

#### Muscle Function Improved Following Either LIV or ZA Treatment

Peak percentage change in grip strength was observed 4 weeks following sham-OVX in the SH mice (Fig.3A). Although all groups exhibited increased grip strength throughout the initial 2 weeks after surgery (OVX), OVX/AI+LIV and OVX/AI+LIV+ZA mice exhibited the sharpest increase among the groups, peaking at an average 125 g of force (Fig.3A), but these were not significant. At the initial 2 weeks after surgery, OVX/AI+LIV mice had greater grip strength (p < 0.05) than OVX/AI mice, and OVX/AI+LIV+ZA mice trended toward increased grip strength (p = 0.06) relative to OVX/AI mice (Fig.3A). At 5 weeks after OVX, grip strength began to decline in all groups (Fig.3A). By week-9, OVX/AI+LIV mice exhibited greater grip strength (p < 0.05) than did OVX/AI mice (Fig.3A). This margin increased (p < 0.007) by week-11, at which point OVX/AI+LIV+ZA mice also exhibited increased grip strength (p < 0.03) relative to OVX/AI mice (Fig.3A). Grip strength of the OVX/AI+LIV mice remained consistently higher than all groups throughout the course of the study following E_2_ deprivation (Fig.3A). Throughout the post-OVX period, an effect (p < 0.01) was observed between groups (Fig.3A).

**Figure 3:**
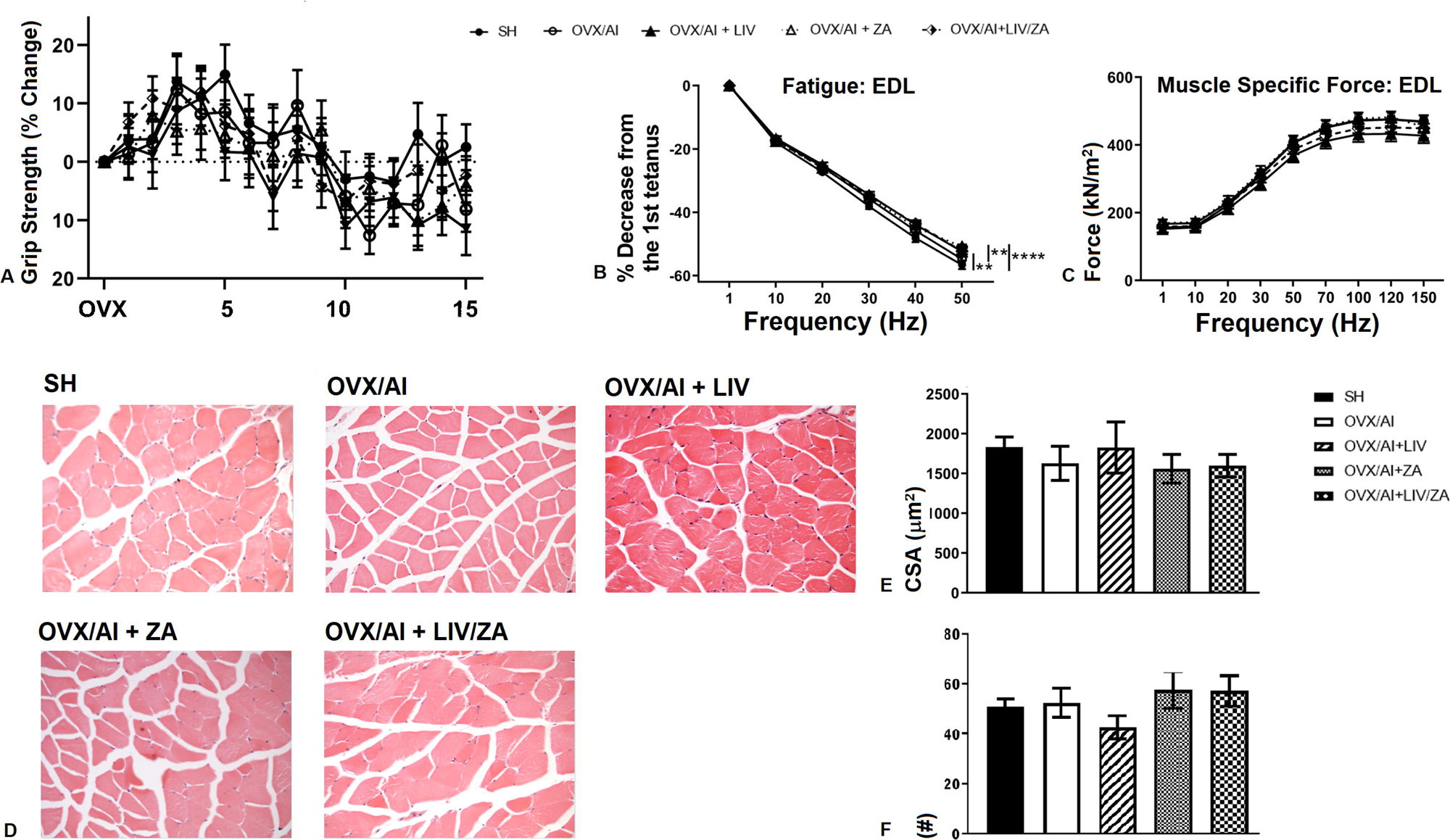
Forelimb grip strength, muscle contractility, and myofiber histology in experiment II. (**A.**) As a measure of muscle function, forelimb grip strength was quantified on a weekly basis. No differences were observed between groups; however, OVX/AI+LIV+ZA mice generally retained the highest grip strength throughout the study. *Ex vivo* muscle contractility force testing was performed at euthanasia on extensor digitorum longus muscles resected from the hindlimbs. Through tetanic stimulation, (**B.**) fatigue and (**C.**) maximum muscle specific force were determined. Muscle fatigue was greatly reduced in both LIV-treated groups at 40 and 50Hz stimulations compared to OVX/AI mice, as did OVX/AI+ZA at the 50Hz stimulation. No differences were observed in specific force in experiment II, (**D.**) Histologic sections (magnification 10×; hematoxylin and eosin) of quadratus femorum were analyzed to quantify average myofiber cross-sectional area and number of myofibers. No significant differences between groups were observed in (**E.**) cross-sectional area or (**F.**) number of myofibers. However, OVX/AI+LIV mice generally had reduced myofiber cross-sectional area. Visually, empty spaces between muscle fibers were evident in untreated OVX/AI mice, and these were nearly absent in sections from the OVX/AI+LIV mice. Abbreviations: OVX, ovariectomy; AI, aromatase inhibitor; LIV, low-intensity vibration; SH, sham-OVX and saline vehicle (control); ZA, zoledronic acid. *p < 0.05; **p < 0.01; ****p < 0.0001.

Percentage decrease from the first tetanus (30-Hz stimulation) during *ex vivo* muscle contractility of the EDL yielded significant improvement in fatigue in OVX/AI+LIV mice (p < 0.03) and OVX/AI+LIV+ZA mice (p < 0.05) relative to SH mice (Fig.3B). At the next frequency (40 Hz) in the stimulation protocol, OVX/AI+LIV and OVX/AI+LIV+ZA mice had less muscle fatigue (p < 0.001 and p < 0.002, respectively) than SH mice (Fig.3B). Additionally, OVX/AI+ZA mice had reduced muscle fatigue (p < 0.05) in contrast to SH mice; however, none of the groups differed from OVX/AI mice (Fig.3B). At 50-Hz stimulation, treatment with LIV and LIV+ZA in OVX/AI mice both led to less muscle fatigue (p < 0.01) than that seen in SH mice, whereas OVX/AI+ZA mice had 10% more muscle fatigue (p < 0.0001) than that observed in SH mice (Fig.3B). At this stimulation frequency, OVX/AI+ZA mice had less muscle fatigue (p < 0.02) than did untreated OVX/AI mice (Fig.3B). No differences were observed in muscle specific force at any of the frequencies used to stimulate the muscle (Fig.3C). Untreated OVX/AI mice showed lower myofiber CSA compared with SH mice (Fig.3D); however, no significant differences in myofiber CSA or number of myofibers were observed across the groups in the skeletally mature mice (Fig.3E and 3F, respectively). Neither OVX/AI+ZA nor OVX/AI+LIV+ZA mice exhibited improvements from OVX/AI vehicle controls.

#### Estrogen Deprivation Increased Perigonadal and Bone Marrow Adipose Accrual, Whereas Either LIV or ZA Alone Reduced Body Weight and Decreased Adiposity

DXA analysis indicated significant increases in % fat mass of OVX/AI mice (p < 0.01) and OVX/AI+ZA mice (p < 0.02) relative to SH mice at 9 weeks after OVX (Fig.4A). By 14 weeks after OVX, OVX/AI mice had increased (p < 0.0001) percent fat mass relative to SH mice; however, LIV treatment alone or in combination did not significantly alter percent fat mass compared to estrogen deprived mice (Fig.4A). No differences were observed in percent lean mass between groups; however, a significant effect (p < 0.0001) was found in all groups with respect to time over the course of the study. Throughout experiment II (older cohort), body weight increased the least in SH mice, and OVX/AI mice had significantly greater (p < 0.01) weight than SH mice. OVX/AI+LIV and OVX/AI+LIV+ZA mice exhibited greater (p < 0.0001) body weight than SH mice by 20 weeks after OVX. Body weight of OVX/AI+ZA mice was lower (p < 0.05) than that of OVX/AI+LIV+ZA mice by the same endpoint. Taken together, these results indicated that young mice treated with LIV had a significantly slower rate of weight and fat gain compared with their untreated counterparts, but older E_2_-deprived mice gained significantly more weight than did their E_2_-replete, SH counterparts.

**Figure 4:**
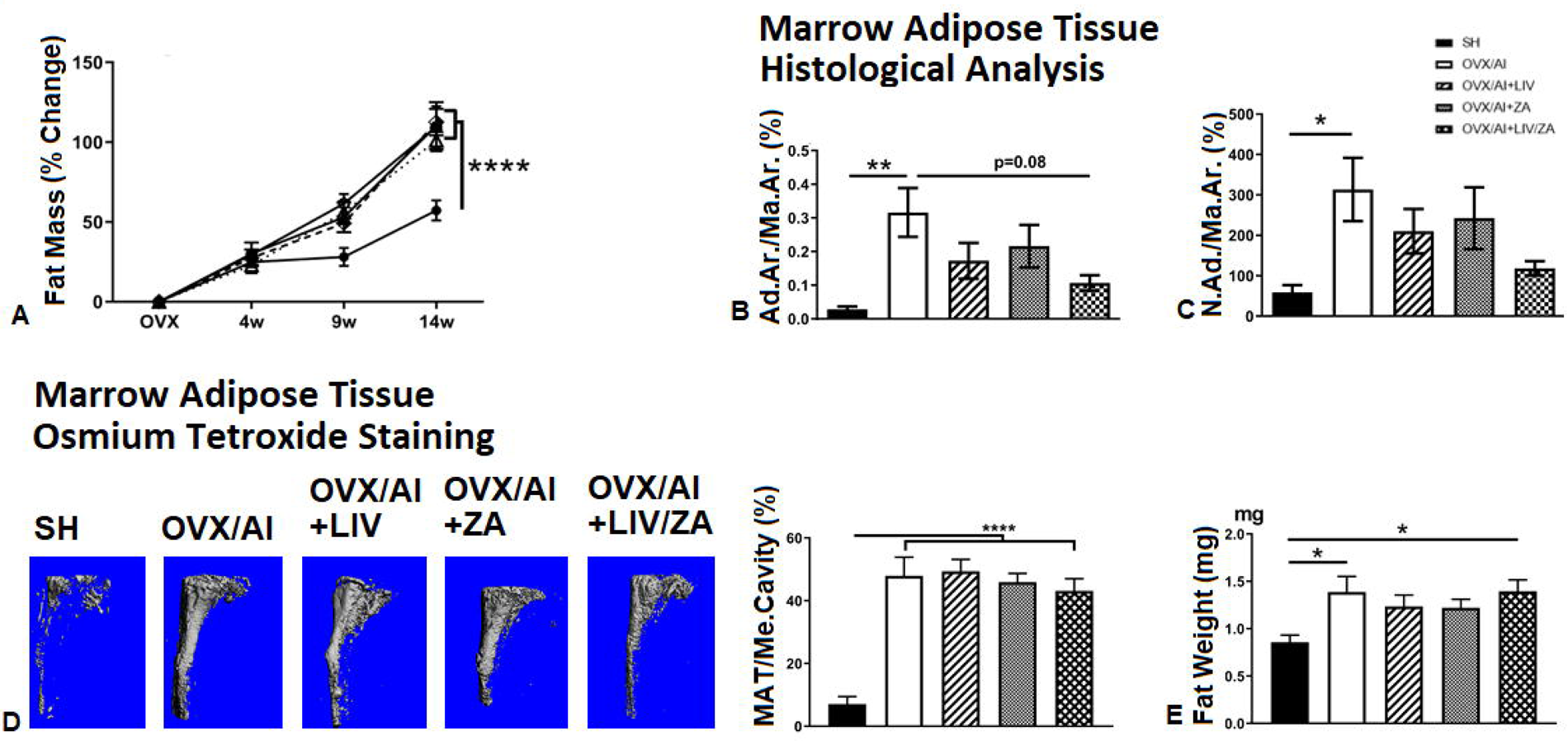
Effect of estrogen deprivation on fat metabolism, with and without treatment with LIV and/or ZA. (**A.**) Total % fat mass measured by dual-energy X-ray absorptiometry (DXA) was significantly increased in all E_2_-deprived mice when compared to SH controls at 14w. Histologic evaluation of (**B.**) Ad.Ar./Ma.Ar. and (**C.**) N.Ad./Ma.Ar. showed significant increases in untreated OVX/AI mice compared with SH mice. All treatment groups had suppressed size and amount of adipose accumulation compared with untreated groups, with OVX/AI+LIV+ZA mice showing the greatest reduction from levels observed in untreated OVX/AI mice. (**D.**) Osmium-tetroxide staining showed significant increases in MAT in all groups that underwent E_2_ deprivation. Visual examination revealed a clear distinction between the tibiae of LIV-treated mice compared with vehicle-treated OVX/AI mice. (**E.**) OVX/AI mice exhibited greater fat pad weights as compared with SH-mice, yet these differences were not significant, possibly due to aging where differences cannot be resolved at that site. Generally, mice treated with LIV that underwent OVX and AI-treatment, however, had lower fad pad mass than those of the OVX/AI mice not treated with LIV. ZA may have had similar trends, though not to the same degree, were observed in either ZA-treated group. Abbreviations: OVX, ovariectomy; AI, aromatase inhibitor; LIV, low-intensity vibration; SH, sham-OVX and saline vehicle (control); ZA, zoledronic acid; Ad.Ar./Ma.Ar., total adipocyte volume encased within a given marrow volume; N.Ad./Ma.Ar., total number of adipocytes within a given marrow volume; Me.Cavity, medullary cavity volume. *p < 0.05; **p < 0.01; ***p < 0.001; ****p < 0.0001.

Ad.Ar./Ma.Ar. in the tibial metaphysis were increased (p < 0.01) in OVX/AI mice relative to SH mice, and these levels were decreased by 15% in OVX/AI+LIV mice (Fig. 4B). Ad.Ar./Ma.Ar. was decreased in OVX/AI+ZA mice by 10% relative to OVX/AI mice, and a reduction was also observed in OVX/AI+LIV+ZA mice (p = 0.08; Fig.4B). N.Ad./Ma.Ar. was increased (p < 0.04) in OVX/AI mice relative to SH mice (Fig.4C). Treatment with LIV resulted in a 33% reduction compared with OVX/AI mice, while N.Ad./Ma.Ar. in OVX/AI+ZA mice decreased by 26%. However, a 62% reduction was observed in OVX/AI+LIV+ZA mice as compared to OVX/AI mice (Fig.4C).

Marrow adipose tissue normalized to medullary cavity volume as quantified by microCT was greater (p < 0.0001) in OVX/AI mice than in SH mice (Fig.4D). OVX/AI+LIV mouse tibiae were not significantly different from those of OVX/AI mice, nor were those of OVX/AI+ZA mice. Marrow adipose tissue decreased in OVX/AI+LIV+ZA mice relative to OVX/AI mice, by 36% compared with SH mice (Fig.4D), but this was not significant.

At euthanasia, both OVX/AI and OVX/AI+LIV+ZA mice had significantly more gonadal fat pad tissue (p < 0.02) than did SH mice (Fig.4E). OVX/AI+LIV mice had more gonadal fat pad tissue than SH mice, but fat pad tissue did not differ between OVX/AI-LIV mice and untreated OVX/AI mice (Fig.4E). None of the treated groups in experiment II significantly differed from OVX/AI mice in terms of gonadal fat pad tissue weight (Fig.4E).

## Discussion

Estrogens serve a central regulatory role in bone remodeling in order to maintain bone health. Aging coupled with menopause contributes to a natural decline in E_2_ synthesis, causing dysregulation in bone remodeling while leading to increased fat and an accompanying decline in muscle strength. Cancer and its many treatment strategies either directly or secondarily disrupt E_2_ homeostasis, and, when combined with the effects of aging and menopause, offsets bone remodeling to favor bone loss while also upregulating the production of fat, which in turn heightens inflammation and crowds out bone remodeling cells in the marrow. Our group has shown that patients with breast cancer bone metastases have both bone loss and muscle weakness attributed to release of sequestered TGF-β from the bone matrix [27]. Therefore, limiting bone loss is critical to the stabilization of disease progression and improving patient outcomes. Our overarching objectives in this study were to quantify the effects of applying a low-magnitude, high-frequency vibratory signal to the whole body as a countermeasure to the negative effects of complete E_2_-deprivation therapy and to determine whether, when coupled with a bisphosphonate (e.g., ZA), LIV could complement the current standard of care for patients diagnosed with breast cancer bone metastases.

E_2_-receptor-positive breast cancers progress through a feed-forward pathway whereby estrogens synthesized from the follicle-stimulating cells of the ovary bind to receptors on tumorigenic cells, driving division. Transformed tumor cells enter the circulation and then have the propensity to seed metastases in the brain, liver, kidney, and bone, inducing osteolytic resorption pits at the endosteal surface in bones. This mechanism is analogous to the androgen-driven effects on tumor progression in prostate cancers, where bone metastases predominantly manifest as sclerotic lesions. Secondary E_2_ sources derive from androgens synthesized by adrenal glands and converted into β-estradiol through aromatase secreted by adipocytes housed in the viscera and the liver. Thus, the use of AI therapy in postmenopausal patients induces complete E_2_-deprivation, significantly decreasing patient mortality by eliminating E_2_, a major accelerator of E_2_-receptor-positive breast cancer progression. Despite improvements in tumor-associated morbidity, E_2_-deprivation exacerbates the deleterious effects of aging and menopause on bone, muscle, and body composition in this patient population. Increased bone fragility is observed in breast cancer patients in remission undergoing AI therapy, raising patient susceptibility to developing fractures, with the highest incidences occurring at the femoral neck and across the lumbar spine [70, 71]. Moreover, the efficacy of long-term anti-hormone treatments in preventing cancer from recurring often wane with time as patients experience increased resistance to treatment [72].

Mechanical signals are fundamentally inseparable from bone maintenance, a reliance perhaps most evident in the setting of physiological unloading (e.g., chronic bedrest, disuse), a condition that aggressively upregulates osteoclast-mediated bone resorption. In patients, age-related osteopenia is differentiated from the extensive cortical lesions derived from bone metastases. Clinical treatment to inhibit osteolysis is achieved through bisphosphonate therapy, but rarely do these lesions heal. Simulated unloading in murine models (e.g., hindlimb suspension models) induces bone catabolism [73] and muscle atrophy [74], but these effects have been reversed through administration of LIV [75]. The vertebrae has been a target for LIV in ameliorating intervertebral disc degeneration [76] in rats with simulated hindlimb unloading. Furthermore, clinical studies have demonstrated that daily administration of LIV can restore the bone mass lost during simulated periods of extended bedrest and improve z-scores for childhood cancer survivors [69]. Additionally, our previous studies showed that treatment of mice bearing spontaneous granulosa cell ovarian cancer [57] and myeloma [58] resulted in suppression of bone loss induced by both endogenous and aggressive cancers.

As anticipated, ZA significantly improved trabecular bone mass and mechanical integrity in the trabecular-rich lumbar vertebral body (L4 and L5) when challenged by the OVX/AI phenotype. However, the effects of ZA were significantly enhanced to levels of E_2_-replete mice when the treatment was complemented with LIV, improving trabecular bone across the lumbar spine, whereas treatment with LIV alone did not impart any benefit in retaining cancellous bone of the appendicular skeleton. Improvements in trabecular bone quantity coincided with improved resistance to fracture from compression testing. These effects on bone, however, were not isolated to cancellous bone, because the cortical bone improved in terms of both quantity and quality (i.e., mechanical strength). What is of specific interest is the greater effect observed when the LIV regimen was combined with ZA, resulting in an outcome that was more effective than ZA alone and which persisted throughout the entirety of the study. We speculate that the mechanism is driven by ZA-induced suppression of osteoclast activity.

Recently, a great deal of attention has focused on the crosstalk between bone and muscle, recognizing that dysfunction in one tissue likely translates to the other. With regards to the present study, compounding the negative effects of AI and breast cancer metastases in swaying bone remodeling toward heightened focal resorption, patients experience reduced muscle strength [77] and elevated body fat [77]. Heightened bone-derived TGF-β, in addition to its autocrine feedback function in metastatic growth, enters the circulation and perpetuates RyR dysfunction and disrupts Ca^2+^ signaling, inducing muscle weakness. Our murine model highlights the accelerated and near-systemic loss of bone associated with E_2_-deprivation. Concomitant loss of lean mass, myofiber number, and myofiber CSA suggest a morphologic relationship between bone and muscle. Histologic sections quantified from the quadratus femora in the younger mouse cohort showed that incorporating daily mechanical loads for an extended period led to muscle fiber hypertrophy in the CSA of hindlimb muscles. These findings may be due to reduced fibrosis or adipocyte deposition, and the increased spacing observed between muscle fibers may stem from reduced oxidative capacity of the muscle fibers or decreased satellite cell maintenance of muscle. Whether the increase in myofiber area is a byproduct of muscle preservation following E_2_ loss or an anabolic response derived from LIV is not fully understood.

Expanding on the functional outcomes, the EDL fatigability observed in OVX/AI mice was similar to the amount detected in SH mice. The inclusion of LIV, regardless of ZA administration, lessened fatigue in the treated mice relative to the muscle fatigue observed in SH mice. However, treatment with ZA alone significantly reduced fatigue in the EDL relative to the values observed in SH mice, but not relative to mice treated with LIV. The reduced fatigue observed in OVX/AI+LIV+ZA mice provides evidence supporting these results. Despite these findings, fatigue in the EDL was unchanged in younger mice exposed to LIV once per day. Reduced EDL fatigue following treatment with LIV+ZA in the older mice suggests that inhibition of skeletal resorption and subtle exposure to LIV elicited a greater response in older mice than in the younger mice. Perhaps the capacity of ZA to prevent TGF-β release contributed to the protection of Ca^2+^ signaling in the muscle. Alternatively, the second bout of daily treatment with LIV could have served to bolster the signal that was only mildly effective in the younger cohort (treated only once per day). The increased grip strength observed in the younger mice supports the clinical observation that mechanical loading (e.g., exercise, physical activity) builds stronger bone and muscle if introduced early in skeletal development. Together, these LIV effects on muscle, either alone or in tandem with ZA, appear to exert multiple effects on composition and functionality. The varied efficacy of the treatment may stem from recruitment of different muscle groups and be a consequence of differing ages. The possible extension of these findings into other muscles, suggested by the increased grip strength observed throughout the study, is of interest.

Bone loss ensuing from E_2_-deprivation therapy illustrates the reliance of bone homeostasis on E_2_ availability while also highlighting the reciprocal relationship between bone and fat. The association of obesity with cancer is gaining more traction [78]; excess fatty tissue is highly correlated with breast cancer incidence [79]. An adipose-enriched phenotype, as paralleled by an obese phenotype, has been associated with a list of maladies, including a body habitus permissive to breast cancers [80–82]. As a consequence of age and diminished E_2_-synthesis, postmenopausal women (and aging men) accumulate fat in visceral tissues and marrow cavities. β-oxidation of excess fatty acids (lipolysis), a secondary means of ATP production, disrupts normal glycolysis in the mitochondria of insulin-sensitive tissues, including muscle and liver [83]. Intermediate metabolites of different carbon chain lengths derived from incomplete transformation are termed acylcarnitines, and their accumulation is associated with insulin resistance [84, 85]. Age-related increases in adiposity are exacerbated by AI therapy, causing adipocyte hypertrophy and infiltration of muscle fibers, displacement of healthy marrow, encasement of critical organs, inflammation, and occlusion of vasculature.

We observed significantly increased gonadal fat deposition in the older mice in response to OVX and daily AI. Additionally, tibial marrow adipose tissue reflected similar increases in adipocyte density and hypertrophy in response to complete E_2_-deprivation. Treatment with LIV drastically reduced both the total area and number of marrow adipocytes and, to a lesser degree, we observed the same effect following treatment with ZA. These results would translate to clinical standard-of-care including anti-resorptive drugs and, if paired with LIV, these drugs could significantly reduce the adiposity (and adverse localized activity) wrought by disease and primary treatments. Therefore, suppressing adiposity in various depots could reduce inflammation and inhibit bone resorption, consistent with upregulation in the lipid profile exhibited by postmenopausal patient cohort being treated with AIs in a previous study [86].

Quantitative serum acylcarnitine analysis identified significant increases in short-chain (C2-C7) acylcarnitine intermediates, specifically acetylcarnitine (C2) and hexanoylcarnitine (C6), in the mice in our study, and these findings served as biomarkers of E_2_-deprivation–related muscle and bone disease, as well as their response to treatment with LIV. Acetylcarnitine has been proposed as a biomarker of hepatocellular carcinoma [87]. Age-dependent increases in acetylcarnitine in mouse and human muscle have been reported [88]. Significant increases in circulating hexanoylcarnitine were identified in response to a 12-week low-calorie diet to induce weight reduction, along with increases in medium- and long-chain acylcarnitines and free fatty acids [89]. The specific source of these acylcarnitines appears to be from fat deposits; however, the specificity of acetylcarnitine and hexanoylcarnitine is not known. Our findings support possible benefits from LIV by suppressing systemic fat mass loss and across various deposits. Peroxisome-proliferator alpha receptor-gamma (PPARγ), a transcriptional regulator of adipocyte differentiation [90], also regulates acylcarnitine metabolism [91]. Previous evidence suggests that LIV directs MSC differentiation towards RunX2-directed pathways and away from PPARγ [60, 92] and induces fat reduction in models of metabolic dysfunction [93, 94]. This is consistent with the data observed in our OVX/AI+LIV mice, suggesting a possible mechanistic connection to LIV-mediated reduction in adiposity and serum acylcarnitines. However, no differences in fat pad weights were observed between all E_2_-deprived mice in the older mouse cohort; therefore, metabolomics was not performed in this cohort, as it did not track with E_2_-deprivation.

The complex mechanisms responsible for driving LIV effects on adiposity may lie, in part, in the ability of mechanical loading to drive the differentiation of the mesenchymal stem cell towards an osteogenic phenotype, thus deterring adipogenic differentiation [95]. To what degree this affects the synthesis of the fatty acids of different classes and across critical tissues remains to be addressed in future experiments. However, the reduction in bone marrow adiposity in the long bones of mice undergoing an extreme and acute deficit in E_2_ without the aid of pharmacologic intervention is a promising finding.

Combining LIV with ZA significantly improved lumbar trabecular bone volume fraction and cortical bone CSA in the femur above either treatment alone, a finding that could be of specific benefit to patients undergoing similar treatments, given that femoral head and vertebral compression fractures are common in patients undergoing anti-E_2_ therapy. Patients who experience fractures have increased recovery times and morbidity. LIV is advantageous as a potential additional arm of treatment, because it does not interrupt other therapeutic efforts. In addition to improved measures of bone, a significant reduction in the fatigability of the EDL via muscle contractility testing was observed in mice jointly treated with LIV and ZA. In short, the benefits to bone were accompanied by improvements in muscle phenotype and strength.

The current study has some limitations deriving from inherent differences in modeling of E_2_-deprivation and the route of LIV administration in mice compared with humans. E_2_-deprivation results in substantial reduction in trabecular bone across the proximal tibiae and distal femur in mice, with mild traces remaining, necessitating the quantitation of the trabecular-rich vertebral bodies of the lumbar spine. Muscle strength was measured with a grip meter using tail suspension over a grip mesh, a method that recruits different muscle groups than a grip test performed in humans, which is quantified exclusively via forelimb grip. Furthermore, that EDL muscle contractility did not improve following either treatment may suggest muscle fiber type and/or location specificity for treatment-induced effects. Perhaps the most important consideration in the study design is the degree of daily handling stress endured by the mice. Despite this, all mice experienced similar handling and subsequently similar downstream stress effects between groups. Additionally, the mechanical signal used in these studies was administered with a singular defined set of parameters. Thus, by varying spatiotemporal parameters, differing effects on the target tissues may be observed. Extending or limiting treatment duration, modulating the waveform frequency or magnitude, or changing the number of treatments could improve outcomes even further. Lastly, the dose of letrozole was determined to be adequate to limit E_2_ synthesis, but these effects, initially observed in younger mice, may not translate completely to skeletally mature and/or aging mice further burdened by accumulating visceral adiposity and waning resident marrow-bound progenitors.

Our study demonstrates that LIV can protect and/or increase lean mass, reduce fat mass, and improve bone endpoints as otherwise compromised by complete E_2_-deprivation, and when LIV is administered twice daily in combination with zoledronic acid, these effects are additive in preserving vertebral trabecular bone. Combining LIV with ZA is unique in the benefit imparted on multiple systems at once. Inhibition of bone resorption with ZA combined with increased bone formation following LIV may aid in patient recovery after treatment or be considered as a pre-surgical strategy to improve patient outcomes for those undergoing anti-hormone treatment.

## Conclusion

In conclusion, our study quantified the negative effects of E_2_-deprivation, a therapeutic modality employed to stop the progression of breast cancer in E_2_-receptor-positive patients, on bone and muscle, while demonstrating the potential of utilizing mechanical signals in the form of LIV and the anti-resorptive drug ZA to inhibit the degree of musculoskeletal deterioration. These findings highlight the combined deleterious impact of diminished E_2_-synthesis and anti-E_2_ treatments, modeled via OVX and AI, in decreasing bone mass and muscle strength and increasing fat due to the absence of circulating E_2_. Although ZA inhibited bone loss, our findings demonstrate the added positive effects of combining low-magnitude mechanical signals with ZA on enhancing preservation and/or increasing lean mass while reducing fat mass, and most impressively, improving trabecular and cortical bone endpoints. Additionally, we identified two novel biomarkers (acetylcarnitine and hexanoylcarnitine) that are increased with E_2_-deprivation-associated musculoskeletal defects and decreased with a successful clinical intervention (e.g., LIV). Under these conditions, our data highlight the benefits of exercise, but more specifically, mechanical loads induced by LIV, in protecting muscle and skeletal tissue and provide the basis to direct future clinical therapies towards combinatorial strategies that incorporate both pharmacologic and non-pharmacologic treatments (Supplementary Fig.3).

## Materials and Methods

### Animal Model

C57BL6 mice (Envigo, Indianapolis, IN, USA) were monitored daily and maintained in accordance with the Institutional Animal Care and Use Committee guidelines at Indiana University and the National Institutes of Health Guidelines for the Care and Use of Laboratory Animals. Mice were housed five per cage at standard room temperature and allowed *ad libitum* access to standard diet-chow (catalog #2018SX, Envigo) and water. A period of 1 week was allotted at the beginning of the study to acclimate the mice to the handling procedures and LIV and mock-LIV treatment protocol. Group designations are detailed in Supplementary Table 1. For experiment I, female 4-week-old C57BL6 mice were divided into two groups (n = 10/group): ovariectomized (OVX) and treated with AI only [OVX/AI(y)] and OVX and treated with AI and LIV [OVX/AI+LIV(y)]. For experiment II, female 15-week-old C57BL6 mice were divided into five groups (n = 20/group): sham-ovariectomized age-matched controls given saline vehicle (SH); OVX and treated with AI only (OVX/AI); OVX and treated with AI and LIV (OVX/AI+LIV); OVX and treated with AI and ZA (OVX/AI+ZA); and OVX and treated with AI, LIV, and ZA (OVX/AI+LIV+ZA).

### Complete E_2_ Deprivation

Sham-surgery and vehicle (phosphate-buffered saline) injection were performed on SH mice, and all other treatment groups underwent OVX supplemented with subcutaneous injection of AI letrozole (Novartis International AG, Basel, Switzerland) administered at a dose of 5 µg/kg per day 5 times per week. Serum β-estradiol was significantly decreased (p < 0.004) in 28-week-old OVX/AI(y) mice and (p < 0.0002) in 28-week-old OVX/AI+LIV(y) mice relative to 4-week-old mice at baseline. No differences in serum β-estradiol were detected between OVX/AI(y) and OVX/AI+LIV(y) mice at 28 weeks of age. This model, characterized in experiment I, was used in a similar fashion for experiment II.

### Bisphosphonate Administration

Low-dose (2.5 ng/kg) ZA (Zometa, Novartis) was subcutaneously injected in the older mice (OVX/AI+ZA and OVX/AI+LIV+ZA groups) once per week commencing the day following OVX surgery.

### Mechanical Loading Protocol

Mechanical loading (LIV and mock-LIV) regimens commenced at 4 weeks of age for the younger cohort (experiment I) and at 16 weeks of age for the older cohort (experiment II). Mice assigned to the mechanical loading regimen were subjected to LIV (0.7 ± 0.025 g, 90 Hz sine wave) for 20 minutes per treatment, administered once per day, 5 days per week over 28 weeks (experiment I), or twice per day, 5 days per week over 24 weeks (experiment II). OVX/AI(y) mice underwent identical handling and loading protocols as OVX/AI+LIV(y) mice but without activation of the platform (mock-LIV). Daily loading consisted of placing mice into individual compartments within a larger container placed on a vertically oscillating platform (modified from BTT Health, Germany) to administer the LIV signal. Displacements required to produce accelerations at 90 Hz induce membrane deformations of approximately 5 microstrain. The lead investigator was not blinded to the experimental groups during the LIV treatment windows.

### Dual-Energy X-ray Absorptiometry (DXA)

Longitudinal *in vivo* measurements of bone mineral density (BMD, g/cm^2^), % whole body fat mass (grams), and % whole body lean mass (grams) were collected via DXA scanning (Piximus 2, Lunar Corporation, Madison, WI, USA) to determine body composition. Prior to each scan, mice were anesthetized with an intraperitoneal injection of ketamine/xylene cocktail (100 mg/kg + 10 mg/kg). Whole body regions of interest excluded the head and tail. DXA analysis of the lumbar spine was performed using a region of interest (60 µm × 20 µm) situated above the sacral spine and extending superior to L1.

### Tissue Harvest and Preservation

At the termination of the study, each mouse was anesthetized using isoflurane inhalation. Euthanasia was confirmed by cervical dislocation. Whole blood collected via cardiac puncture was stored in serum-separation tubes, with serum aliquots stored at −80°C. Bones for histologic processing, including the right femora and tibiae, were stripped of muscle and residual soft tissue and fixed in 10% neutral buffered formalin, which was replaced at 48 hours with 70% ethanol. At euthanasia, lumbar vertebrae and femora harvested for mechanical testing were wrapped in saline-soaked gauze and frozen at −20°C. The quadratus femoris muscle was formalin-fixed for 48 hours following euthanasia and then transferred to 70% ethanol and preserved for downstream processing.

### Micro-Computed Tomography (MicroCT)

*In vivo* and *ex vivo* scans were performed to quantify bone microarchitecture and visualize reconstructed 3-dimensional representations. Distal femora and L4 and L5 vertebrae were measured *ex vivo* using X-ray microCT (μCT40, Scanco Medical, Wayne, PA, USA). X-ray parameters were set to the following: *in vivo* source voltage E = 55 kVp, current = 145 μA, integration time = 300 ms, and voxel size = 19 µm; *ex vivo* source voltage E = 55 kVp, current = 145 μA, integration time = 300 ms, and voxel size = 10 μm. Cancellous (trabecular) bone parameters were identified using the following nomenclature: Tb.BV/TV, Tb.Conn.D, Tb.N, trabecular thickness (Tb.Th), trabecular separation (Tb.Sp), trabecular BMD (Tb.BMD), and trabecular structure model index (Tb.SMI). Cortical bone parameters were identified using the following nomenclature: cortical bone area fraction (Ct.BA/TA), cortical thickness (Ct.Th), and cortical polar moment of inertia (Ct.pMOI). Starting 700 μm proximal to the femoral growth plate, 1000 μm of metaphyseal trabecular bone was evaluated. A 500-μm uniform length of trabecular bone (excluding the growth plate) was evaluated across the L5 and L4 vertebral bodies; 500 µm of cortical bone was quantified about the femoral mid-diaphysis.

### Mechanical Testing of Bones

Mechanical testing was performed using a 3-point bending jig to quantify both structural and material level cortical bone properties at the femoral mid-diaphysis, and vertical compression jigs were used to quantify the structural properties resisting fracture in the lumbar vertebrae. Force and displacement data were recorded (Rcontroller, Test Resources, Shakopee, MN, USA) and structural and material properties were derived from force curves using a customized Matlab (MathWorks, Natick, MA, USA) script.

Spines were thawed; then L4 and L5 vertebrae were isolated using a scalpel and cleaned of intervertebral discs and cartilaginous tissue from proximal and distal ends. Pedicles and spinous processes were removed, leaving only the vertebral body. Sample vertebrae were loaded along the vertical axis with 2N of preload force between platens using 11 lb-f load cells. Position-controlled rate of compression was applied until failure. Samples that slipped during compression were omitted from the dataset.

Femora were thawed, excess soft tissue was removed, and the femora were scanned using microCT to obtain CSA. For 3-point mechanical testing of the femora, the distance between the lower supports (Δx = 9 mm) was maintained throughout the analyses while the upper support was centered between the bottom supports. 150 lb-f load cells were used for testing with 1N of preload force, and the position-controlled rate was applied to each sample (femora were positioned anterior side facing up) until failure.

### Bone Histology/Histomorphometry

The lengths of long bones were measured *ex vivo* using digital calipers. Formalin-fixed bone specimens for paraffin embedding and osmium tetroxide staining were decalcified in ethylenediaminetetraacetic acid (10%; pH = 7.32) for 2 weeks. 5 μm paraffin-embedded sagittal cross sections were stained with hematoxylin and eosin. PMMA-embedded intact lumbar vertebrae were sectioned (4.5 µm) and were subsequently either left unstained (i.e., dynamic histomorphometry) or stained with tartrate-resistant acid phosphatase to quantify osteoclasts or with von Kossa-McNeil to quantify osteoblasts. Bone analysis software (v.18.1, BIOQUANT Osteo, Image Analysis Corporation, Nashville, TN, USA) was used to analyze the results.

### Forelimb Grip Strength

Muscle function was quantified with the use of a grip strength meter (Bioseb, Pinellas Park, FL, USA) to report maximum applied force (grams). Prior to each pass the researcher suspended mice individually by their tails, placing forelimbs within reaching distance of the front of the mesh grid. A constant horizontal pulling force was applied until the mice released their grip from the mesh. Triplicate readings per mouse were averaged and longitudinal measurements were taken throughout the study.

### Muscle Histology

Paraffin-embedded histologic sections (4.5 µm) of quadratus femorum were stained with hematoxylin and eosin. Dedicated software (BIOQUANT Osteo Image Analysis Corporation) was used to acquire 5 fields of view at 10x magnification and then used to quantify myofiber CSA by contouring each myofiber within the field of view, thresholding out the space between the fibers, and normalizing to the total tissue area.

### Muscle Specific Force/Muscle Fatigue

*Ex vivo* contractility of the extensor digitorum longus (EDL) muscle was performed via tetanic stimulation as previously described [27]. Briefly, whole EDL were dissected from hindlimbs and placed in Tyrode buffer to maintain Ca^2+^ potential. Muscles were mounted to force transducers (Aurora Scientific, Ontario, Canada) using stainless steel hooks tied to the proximal and distal tendons, while remaining hydrated in a stimulation chamber containing O_2_/CO_2_ (95/5%) bubbled into Tyrode buffer. Excised muscle was stimulated to contract using a supramaximal stimulus between two platinum electrodes. Force–frequency and fatigue–frequency relationships were determined by triggering contraction using incremental stimulation frequencies (specific force: 0.5 ms pulses at 1–150 Hz for 350 ms at maximal voltage; fatigue: 0.5 ms pulses at 1–50 Hz for 350 ms at maximal voltage). Data were collected using proprietary “Dynamic Muscle Control/Data Acquisition” and “Dynamic Muscle Control Data Analysis” programs (Aurora Scientific). Wet tissue weights and lengths of gastrocnemius, EDL, soleus, and tibialis anterior muscles were recorded with digital calipers at euthanasia. For quantification of the specific force and fatigue of the EDL, the absolute force was normalized to muscle size and calculated by dividing muscle weight by muscle length using a muscle density constant (1.056 kg/m^-3^) [96]. Investigators performing the measurements were blinded to the treatment groups.

### Marrow Adipose Histology

Histologic sections (20×, hematoxylin and eosin) of the distal tibial metaphysis were analyzed starting from the growth plate for 2000 µm distally to quantify the total adipocyte volume encased within a given marrow volume (Ad.Ar./Ma.Ar.) and total number of adipocytes within this space (N.Ad./Ma.Ar.). Briefly, the region of interest across the marrow space was contoured followed by binary thresholding of the marrow and adipocytes independently, which were each quantified using Osteo (v.19.2).

### Osmium-Tetroxide Staining of Marrow Adipose Tissue

Marrow adipose tissue was quantified as described previously [97]. Formalin-fixed tibiae were scanned via microCT. Bones were decalcified in ethylenediaminetetraacetic acid (pH = 7.32) for 10 days, rinsed with tap water, and transferred to Eppendorf tubes. A 1:1 ratio of osmium tetroxide (OsO_4_) to sodium bicarbonate mixture was added to each sample, allowing the OsO_4_ to permeate the marrow for 48 hours. After discarding the waste solution, bones were washed with tap water again and stored in phosphate-buffered saline at 4°C. Samples were scanned using microCT imaging (E = 55 kVa, I = 72 µA, voxel size = 12.5 µm). Values were produced by thresholding tissue that absorbed OsO_4_ and normalizing this value to the medullary cavity volume across 6000 µm from the epiphysis to the inferior tibiofibular joint.

### Immunoassays

Serum was collected at the termination of the study and stored at −80°C. Thawed samples were used to quantify 17β-estradiol (Calbiotech, catalog #ES180S-100, El Cajon, CA, USA) and leptin (Quantikine, catalog #MOB00, Minneapolis, MN, USA) as analyzed by enzyme-linked immunosorbent assay per manufacturer guidelines.

### Free Fatty Acid Serum Metabolite Analyses

Serum isolated from whole blood at baseline and euthanasia was used to quantify free fatty acids. Acylcarnitines were analyzed using stable isotope dilution techniques. Amino acids and acylcarnitine measurements were made by flow injection tandem mass spectrometry using sample preparation methods described previously [98–100]. The data were acquired using an Acquity UPLC system (Waters Corporation, Milford, MA, USA) equipped with a triple quadrupole detector and a data system controlled by MassLynx operating system (v.4.1, Waters Corporation). Organic acids were quantified using methods described previously employing Trace Ultra GC coupled to ISQ MS operating under Xcalibur (v.2.2, Thermo Fisher Scientific, Inc., Waltham, MA, USA). Acylcarnitine concentrations were analyzed using Metaboanalyst (v2.0, McGill, Quebec, Canada) [101–103] and run in the statistical package R (v.2.14.0). Initial unsupervised evaluation using principal component analysis identified the mouse strain as the primary source of variance; therefore, the metabolites were mean-centered per strain after log transformation of concentrations, resulting in a Gaussian distribution. All metabolite analyses were performed blinded to group, experimental treatment, and all other identifying data. After acylcarnitine values were determined, metabolomics analysis of grouped data was performed with the user blinded to all group designations.

### Glucose Tolerance Testing

Mice aged to 26 weeks were fasted overnight and intraperitoneally injected the following morning with a bolus of glucose (20% glucose solution). Subsequent blood glucose samples drawn from the tail vein were read using a glucometer at 0-, 15-, 30-, 60-, and 120-minute intervals. Serum glucose was not measured in the older mice in order to minimize potential stress.

### Statistical Analysis

All graphical data are presented with bars representing means ± standard error. Two-way analysis of variance was performed for longitudinal measurements of body weights, grip strength, DXA analyses, muscle contractility, and microCT followed by Tukey post hoc analysis (Prism, GraphPad Software v.7.0c, San Diego, CA, USA). Pearson correlations were performed using a 95% confidence interval (p < 0.05) for comparing total adipose tissue with serum leptin. The Student *t* test was performed on single-timepoint data. Longitudinal, groupwise comparisons were made between SH, OVX/AI, OVX/AI+LIV, OVX/AI+ZA, and OVX/AI+LIV+ZA by two-way analysis of variance followed by Tukey post hoc analysis. Single-timepoint analyses were quantified by one-way analysis of variance and Tukey post-hoc analyses. A moderated t-statistic model (SAM) [104] was used to identify metabolites that differed between groups at a false discovery rate <5% with Fisher least significant difference used for posttest comparisons. Power calculations accounted for effect size (0.25) and for statistical power ≥ 0.8.

## List of Supplementary Materials

Supplementary Material for this manuscript includes the following:

Supplementary Figure 1: Serum metabolomics analysis or short-chain free fatty acids.

Supplementary Figure 2: Serum metabolomics analysis of medium and long-chain free fatty acids.

Supplementary Figure 3: Depiction of the *in vivo* effects of complete estrogen deprivation in mice treated with aromatase inhibitor letrozole on bone, muscle and fat and how mechanical loading in the form of low intensity vibration combined with zoledronic acid suppress these outcomes.

Table S1: Group designation and treatments administration as applicable administered in the younger cohort of mice (Experiment I) and older cohort of mice (Experiment II).

Table S2: Micro-computed tomography data of vertebral and femoral bone microarchitecture.

Table S3: Mechanical testing data reflecting cortical (3pt-bending) and trabecular (compression testing) bone properties derived from femora and vertebrae, respectively, from experiment II.

## Acknowledgements

The authors would like to express their gratitude to the Indiana University Laboratory Animal Resource Center and to the Matthew Allen Lab personnel at IUSM, specifically, Mohammad W. Aref, Elizabeth A. Swallow, and Corrine E. Metzger, for their assistance during post-processing and access to mechanical testing equipment; and to Erica Goodoff from The University of Texas MD Anderson Cancer Center Research Medical Library.

## Author Contributions

GMP, TT, LEW, KSM, and TAG conceptualized the experiments. GMP, TT, SKJ, and YS provided husbandry, logging, and technical care with the mice. GMP, TT, RRP, SM, KSM, and TAG provided histological and bone histomorphometry expertise and analyses. GMP, SKJ, RRP, and SS performed µCT scanning and analysis. GMP, LEW, and RRP performed MAT staining and analysis. GMP and TT performed glucose metabolism studies and analysis. MW provided technical advice and performed analysis of metabolomics data. GMP, TT, and LEW performed ex vivo analysis of muscle contractility. GMP, TT, SKJ RRP and YS performed and analyzed DXA scans. GMP performed mechanical testing on femora and vertebrae and subsequent analysis. GMP, WRT, and CTR provided expertise on mechanical loading. TT, LEW, KSM, and TAG provided expertise on breast cancer treatment and estrogen deprivation procedures and outcomes. GMP and TAG wrote the manuscript with input from all authors. All authors contributed to the discussion of the results over the course of the experiments and provided critical review of the manuscript.

## Data and Materials Availability

All bone data associated with this study are available in the main text or supplementary materials.

## Supplementary Figures and Tables

**Supplementary Figure 1:** Serum metabolomics analysis showed that short chain acylcarnitines (C2-C7) were significantly elevated in OVX/AI(y) and OVX/AI+LIV(y) mice after 28 weeks of E_2_-deprivation relative to baseline readings. However, OVX/AI+LIV(y) mice also had significantly lower levels of C2-C7 compared with OVX/AI(y)-mice. Abbreviations: OVX, ovariectomy; AI, aromatase inhibitor; LIV, low intensity vibration.

**Supplementary Figure 2:** Serum metabolomics analysis showed an increase in medium-chain free fatty acids but not long-chain free fatty acids.

**Supplementary Figure 3:** Aromatase inhibitor treatment-associated bone loss and muscle weakness arises in patients undergoing aromatase inhibitor therapy, which is used exclusively to inhibit the synthesis of peripheral estrogens. Estrogen deprivation, as in postmenopausal women, also increases the burden of adipose in various compartments. The bisphosphonate zoledronic acid (ZA) is commonly employed in the clinic to suppress osteoclast activity and is known to deter the progression of the tumor upon metastasis to the bone. Formation of new bone or healing of diseased bone is not a significant benefit afforded through the administration of zoledronic acid. Incorporating LIV alone results in the formation of new bone and/or inhibited osteoclast-mediated bone resorption, protecting muscle fiber size, and slows the gain of fat. Using the two treatment strategies in parallel appears to allow for greater trabecular and cortical bone mass, which translates into bones that are more resistant to mechanical failure. LIV has the effect of preserving lean mass, inducing muscle fiber hypertrophy and slowing the gain of fat when challenged with estrogen deprivation.

**Supplemental Table 1:** Table dictating the experimental design of the project. Experiment I consisted of two groups of n = 10 randomly assigned 4-week-old female C57Bl/6 mice/group: ovariectomized (OVX) and treated daily with aromatase inhibitor (AI) letrozole to induce complete E_2_ deprivation, either treated (OVX/AI+LIV(y)) or mock-treated (OVX/AI(y)) with LIV once per day, 5 days per week for 4 weeks, at which point all mice underwent E_2_-deprivation. LIV or mock-LIV continued for an additional 24w following surgery. Experiment II consisted of 5 groups (n = 20/group) of 15-week-old female C57Bl/6 mice: estrogen replete, sham-OVX age-matched controls given saline vehicle (SH); OVX and treated with AI only (OVX/AI); OVX and treated with AI and LIV (OVX/AI+LIV); OVX and treated with AI and ZA (OVX/AI+ZA); and OVX and treated with AI, LIV, and ZA (OVX/AI+LIV+ZA). Mice were treated or mock-treated with LIV for 5 weeks at which point mice underwent either sham surgery or complete E_2_ deprivation, as described in Experiment I, with LIV and mock-LIV treatments continuing thereafter until the study termination. 3 weeks following surgery, mice were treated with ZA or vehicle-control once per week.

**Supplemental Table 2:** *Ex vivo* microCT analyses of L5 vertebrae and femora. Experiment I quantified trabecular (Tb.) parameters across the L5 to highlight significantly greater Tb. bone quantity. Similar analyses were performed in Experiment II, highlighting significant loss of bone in E_2_-deprived mice, both in the L5 as well as the cortical compartment of the femur. Tb. and Ct. parameters improved with the addition of ZA relative to E_2_-deprived controls, and these increased further with the combination of LIV and ZA.

**Supplemental Table 3:** 3-pt bending mechanical testing properties of the femur (Ct.) to derive structural and tissue-level properties. Significant reductions in yield force, ultimate force, and stiffness were observed in the femur in response to E_2_ deprivation without treatment relative to sham controls. Either LIV or ZA treatment alone improved these outcomes relative to E_2_-deprived mice without treatment, but when combined the mechanical resistance to fracture of the femur was improved even further than with either treatment alone. Compression testing of the L5 vertebrae (Tb.) exhibited similar decline in bone following E_2_ deprivation as compared to sham controls, but which was improved considerably with ZA, and even further once LIV was combined with ZA.

## References and Notes

1. Kingsley, L.A., et al., Molecular biology of bone metastasis. Mol Cancer Ther, 2007. 6(10): p. 2609–17.

2. Siegel, R.L., et al., Cancer statistics, 2022. CA Cancer J Clin, 2022. 72(1): p. 7–33.

3. Kohler, B.A., et al., Annual Report to the Nation on the Status of Cancer, 1975-2011, Featuring Incidence of Breast Cancer Subtypes by Race/Ethnicity, Poverty, and State. J Natl Cancer Inst, 2015. 107(6): p. djv048.

4. Clark, G.M., C.K. Osborne, and W.L. McGuire, Correlations between estrogen receptor, progesterone receptor, and patient characteristics in human breast cancer. J Clin Oncol, 1984. 2(10): p. 1102–9.

5. Weilbaecher, K.N., T.A. Guise, and L.K. McCauley, Cancer to bone: a fatal attraction. Nat Rev Cancer, 2011. 11(6): p. 411–25.

6. Tubiana-Hulin, M., Incidence, prevalence and distribution of bone metastases. Bone, 1991. 12 Suppl 1: p. S9–10.

7. Buijs, J.T., K.R. Stayrook, and T.A. Guise, TGF-beta in the Bone Microenvironment: Role in Breast Cancer Metastases. Cancer Microenviron, 2011. 4(3): p. 261–81.

8. Guise, T.A., et al., The combined effect of tumor-produced parathyroid hormone-related protein and transforming growth factor-alpha enhance hypercalcemia in vivo and bone resorption in vitro. J Clin Endocrinol Metab, 1993. 77(1): p. 40–5.

9. Guise, T.A., et al., Evidence for a causal role of parathyroid hormone-related protein in the pathogenesis of human breast cancer-mediated osteolysis. J Clin Invest, 1996. 98(7): p. 1544–9.

10. Guo, D., J. Huang, and J. Gong, Bone morphogenetic protein 4 (BMP4) is required for migration and invasion of breast cancer. Mol Cell Biochem, 2012. 363(1-2): p. 179–90.

11. Mohan, S. and D.J. Baylink, Bone growth factors. Clin Orthop Relat Res, 1991(263): p. 30–48.

12. Javelaud, D., et al., TGF-beta/SMAD/GLI2 signaling axis in cancer progression and metastasis. Cancer Res, 2011. 71(17): p. 5606–10.

13. Roodman, G.D., Mechanisms of bone metastasis. N Engl J Med, 2004. 350(16): p. 1655–64.

14. Love, R.R., et al., Effects of tamoxifen on bone mineral density in postmenopausal women with breast cancer. N Engl J Med, 1992. 326(13): p. 852–6.

15. Fisher, B., et al., A randomized clinical trial evaluating tamoxifen in the treatment of patients with node-negative breast cancer who have estrogen-receptor-positive tumors. N Engl J Med, 1989. 320(8): p. 479–84.

16. Burstein, H.J., et al., American Society of Clinical Oncology clinical practice guideline: update on adjuvant endocrine therapy for women with hormone receptor-positive breast cancer. J Clin Oncol, 2010. 28(23): p. 3784–96.

17. Shiau, A.K., et al., The structural basis of estrogen receptor/coactivator recognition and the antagonism of this interaction by tamoxifen. Cell, 1998. 95(7): p. 927–37.

18. Dowsett, M., et al., Meta-analysis of breast cancer outcomes in adjuvant trials of aromatase inhibitors versus tamoxifen. J Clin Oncol, 2010. 28(3): p. 509–18.

19. Simpson, E.R. and S.R. Davis, Minireview: aromatase and the regulation of estrogen biosynthesis--some new perspectives. Endocrinology, 2001. 142(11): p. 4589–94.

20. Goss, P.E., et al., Exemestane for breast-cancer prevention in postmenopausal women. N Engl J Med, 2011. 364(25): p. 2381–91.

21. Coates, A.S., et al., Five years of letrozole compared with tamoxifen as initial adjuvant therapy for postmenopausal women with endocrine-responsive early breast cancer: update of study BIG 1-98. J Clin Oncol, 2007. 25(5): p. 486–92.

22. Saad, F., et al., Cancer treatment-induced bone loss in breast and prostate cancer. J Clin Oncol, 2008. 26(33): p. 5465–76.

23. Rabaglio, M., et al., Bone fractures among postmenopausal patients with endocrine-responsive early breast cancer treated with 5 years of letrozole or tamoxifen in the BIG 1-98 trial. Ann Oncol, 2009. 20(9): p. 1489–98.

24. Schneider, A., et al., Bone turnover mediates preferential localization of prostate cancer in the skeleton. Endocrinology, 2005. 146(4): p. 1727–36.

25. Bellinger, A.M., et al., Remodeling of ryanodine receptor complex causes “leaky” channels: a molecular mechanism for decreased exercise capacity. Proc Natl Acad Sci U S A, 2008. 105(6): p. 2198–202.

26. Andersson, D.C., et al., Ryanodine receptor oxidation causes intracellular calcium leak and muscle weakness in aging. Cell Metab, 2011. 14(2): p. 196–207.

27. Waning, D.L., et al., Excess TGF-beta mediates muscle weakness associated with bone metastases in mice. Nat Med, 2015. 21(11): p. 1262–1271.

28. McPherson, P.S. and K.P. Campbell, The ryanodine receptor/Ca2+ release channel. J Biol Chem, 1993. 268(19): p. 13765–8.

29. Bekker, P.J., et al., A single-dose placebo-controlled study of AMG 162, a fully human monoclonal antibody to RANKL, in postmenopausal women. 2004. J Bone Miner Res, 2005. 20(12): p. 2275–82.

30. Lipton, A., et al., Randomized active-controlled phase II study of denosumab efficacy and safety in patients with breast cancer-related bone metastases. J Clin Oncol, 2007. 25(28): p. 4431–7.

31. Gnant, M., et al., Adjuvant denosumab in breast cancer (ABCSG-18): a multicentre, randomised, double-blind, placebo-controlled trial. Lancet, 2015. 386(9992): p. 433–43.

32. Ottewell, P.D., et al., Zoledronic acid has differential antitumor activity in the pre- and postmenopausal bone microenvironment in vivo. Clin Cancer Res, 2014. 20(11): p. 2922–32.

33. Hughes, D.E., et al., Bisphosphonates promote apoptosis in murine osteoclasts in vitro and in vivo. J Bone Miner Res, 1995. 10(10): p. 1478–87.

34. Jagdev, S.P., et al., The bisphosphonate, zoledronic acid, induces apoptosis of breast cancer cells: evidence for synergy with paclitaxel. Br J Cancer, 2001. 84(8): p. 1126–34.

35. Senaratne, S.G., et al., Bisphosphonates induce apoptosis in human breast cancer cell lines. Br J Cancer, 2000. 82(8): p. 1459–68.

36. Lintermans, A., et al., Aromatase inhibitor-induced loss of grip strength is body mass index dependent: hypothesis-generating findings for its pathogenesis. Ann Oncol, 2011. 22(8): p. 1763–9.

37. Henry, N.L., J.T. Giles, and V. Stearns, Aromatase inhibitor-associated musculoskeletal symptoms: etiology and strategies for management. Oncology (Williston Park), 2008. 22(12): p. 1401–8.

38. Brown, S.A. and T.A. Guise, Cancer treatment-related bone disease. Crit Rev Eukaryot Gene Expr, 2009. 19(1): p. 47–60.

39. Donnellan, P.P., et al., Aromatase inhibitors and arthralgia. J Clin Oncol, 2001. 19(10): p. 2767.

40. Henry, N.L., et al., Predictors of aromatase inhibitor discontinuation as a result of treatment-emergent symptoms in early-stage breast cancer. J Clin Oncol, 2012. 30(9): p. 936–42.

41. Henry, N.L., et al., Prospective characterization of musculoskeletal symptoms in early stage breast cancer patients treated with aromatase inhibitors. Breast Cancer Res Treat, 2008. 111(2): p. 365–72.

42. Dent, S.F., et al., Aromatase inhibitor therapy: toxicities and management strategies in the treatment of postmenopausal women with hormone-sensitive early breast cancer. Breast Cancer Res Treat, 2011. 126(2): p. 295–310.

43. Stattin, K., et al., Leisure-Time Physical Activity and Risk of Fracture: A Cohort Study of 66,940 Men and Women. J Bone Miner Res, 2017. 32(8): p. 1599–1606.

44. Hinton, P.S., P. Nigh, and J. Thyfault, Effectiveness of resistance training or jumping-exercise to increase bone mineral density in men with low bone mass: A 12-month randomized, clinical trial. Bone, 2015. 79: p. 203–12.

45. Dalsky, G.P., et al., Weight-bearing exercise training and lumbar bone mineral content in postmenopausal women. Ann Intern Med, 1988. 108(6): p. 824–8.

46. Romijn, J.A., et al., Regulation of endogenous fat and carbohydrate metabolism in relation to exercise intensity and duration. Am J Physiol, 1993. 265(3 Pt 1): p. E380–91.

47. Styner, M., et al., Bone marrow fat accumulation accelerated by high fat diet is suppressed by exercise. Bone, 2014. 64: p. 39–46.

48. Nicklas, B.J., et al., Diet-induced weight loss, exercise, and chronic inflammation in older, obese adults: a randomized controlled clinical trial. Am J Clin Nutr, 2004. 79(4): p. 544–51.

49. Huang, H.P., et al., Adherence to prescribed exercise time and intensity declines as the exercise program proceeds: findings from women under treatment for breast cancer. Support Care Cancer, 2015. 23(7): p. 2061–71.

50. Ness, K.K., et al., Skeletal, neuromuscular and fitness impairments among children with newly diagnosed acute lymphoblastic leukemia. Leuk Lymphoma, 2015. 56(4): p. 1004–11.

51. Ness, K.K., et al., Physiologic frailty as a sign of accelerated aging among adult survivors of childhood cancer: a report from the St Jude Lifetime cohort study. J Clin Oncol, 2013. 31(36): p. 4496–503.

52. Fritton, S.P., K.J. McLeod, and C.T. Rubin, Quantifying the strain history of bone: spatial uniformity and self-similarity of low-magnitude strains. J Biomech, 2000. 33(3): p. 317–25.

53. Rubin, C., et al., Anabolism. Low mechanical signals strengthen long bones. Nature, 2001. 412(6847): p. 603–4.

54. Pagnotti, G.M., et al., Combating osteoporosis and obesity with exercise: leveraging cell mechanosensitivity. Nat Rev Endocrinol, 2019. 15(6): p. 339–355.

55. Huang, R.P., C.T. Rubin, and K.J. McLeod, Changes in postural muscle dynamics as a function of age. J Gerontol A Biol Sci Med Sci, 1999. 54(8): p. B352–7.

56. Morse, C.I., et al., In vivo physiological cross-sectional area and specific force are reduced in the gastrocnemius of elderly men. J Appl Physiol (1985), 2005. 99(3): p. 1050–5.

57. Pagnotti, G.M., et al., Low magnitude mechanical signals mitigate osteopenia without compromising longevity in an aged murine model of spontaneous granulosa cell ovarian cancer. Bone, 2012. 51(3): p. 570–7.

58. Pagnotti, G.M., et al., Low intensity vibration mitigates tumor progression and protects bone quantity and quality in a murine model of myeloma. Bone, 2016. 90: p. 69–79.

59. Patel, V.S., et al., Incorporating Refractory Period in Mechanical Stimulation Mitigates Obesity-Induced Adipose Tissue Dysfunction in Adult Mice. Obesity (Silver Spring), 2017. 25(10): p. 1745–1753.

60. Luu, Y.K., et al., Mechanical stimulation of mesenchymal stem cell proliferation and differentiation promotes osteogenesis while preventing dietary-induced obesity. J Bone Miner Res, 2009. 24(1): p. 50–61.

61. Frechette, D.M., et al., Diminished satellite cells and elevated adipogenic gene expression in muscle as caused by ovariectomy are averted by low-magnitude mechanical signals. J Appl Physiol (1985), 2015. 119(1): p. 27–36.

62. Uzer, G., et al., Cell Mechanosensitivity to Extremely Low-Magnitude Signals Is Enabled by a LINCed Nucleus. Stem Cells, 2015. 33(6): p. 2063–76.

63. Uzer, G., et al., Concise Review: Plasma and Nuclear Membranes Convey Mechanical Information to Regulate Mesenchymal Stem Cell Lineage. Stem Cells, 2016. 34(6): p. 1455–63.

64. Sen, B., et al., Mechanical signal influence on mesenchymal stem cell fate is enhanced by incorporation of refractory periods into the loading regimen. J Biomech, 2011. 44(4): p. 593–9.

65. Yi, X., et al., Mechanical suppression of breast cancer cell invasion and paracrine signaling to osteoclasts requires nucleo-cytoskeletal connectivity. Bone Res, 2020. 8(1): p. 40.

66. Gilsanz, V., et al., Low-level, high-frequency mechanical signals enhance musculoskeletal development of young women with low BMD. J Bone Miner Res, 2006. 21(9): p. 1464–74.

67. DiVasta, A.D., et al., The ability of low-magnitude mechanical signals to normalize bone turnover in adolescents hospitalized for anorexia nervosa. Osteoporos Int, 2017. 28(4): p. 1255–1263.

68. Wren, T.A., et al., Effect of high-frequency, low-magnitude vibration on bone and muscle in children with cerebral palsy. J Pediatr Orthop, 2010. 30(7): p. 732–8.

69. Mogil, R.J., et al., Effect of Low-Magnitude, High-Frequency Mechanical Stimulation on BMD Among Young Childhood Cancer Survivors: A Randomized Clinical Trial. JAMA Oncol, 2016. 2(7): p. 908–14.

70. Bouvard, B., et al., High prevalence of vertebral fractures in women with breast cancer starting aromatase inhibitor therapy. Ann Oncol, 2012. 23(5): p. 1151–6.

71. Villa, P., et al., Impact of aromatase inhibitor treatment on vertebral morphology and bone mineral density in postmenopausal women with breast cancer. Menopause, 2016. 23(1): p. 33–9.

72. Ma, C.X., et al., Mechanisms of aromatase inhibitor resistance. Nat Rev Cancer, 2015. 15(5): p. 261–75.

73. Gupta, S., et al., Multiple exposures to unloading decrease bone’s responsivity but compound skeletal losses in C57BL/6 mice. Am J Physiol Regul Integr Comp Physiol, 2012. 303(2): p. R159–67.

74. Lawler, J.M., W. Song, and S.R. Demaree, Hindlimb unloading increases oxidative stress and disrupts antioxidant capacity in skeletal muscle. Free Radic Biol Med, 2003. 35(1): p. 9–16.

75. Ozcivici, E. and S. Judex, Trabecular bone recovers from mechanical unloading primarily by restoring its mechanical function rather than its morphology. Bone, 2014. 67: p. 122–9.

76. Holguin, N., et al., Brief daily exposure to low-intensity vibration mitigates the degradation of the intervertebral disc in a frequency-specific manner. J Appl Physiol (1985), 2011. 111(6): p. 1846–53.

77. Ballinger, T.J., et al., Impact of primary breast cancer therapy on energetic capacity and body composition. Breast Cancer Res Treat, 2018. 172(2): p. 445–452.

78. Quail, D.F., et al., Obesity alters the lung myeloid cell landscape to enhance breast cancer metastasis through IL5 and GM-CSF. Nat Cell Biol, 2017. 19(8): p. 974–987.

79. Iyengar, N.M., et al., Association of Body Fat and Risk of Breast Cancer in Postmenopausal Women With Normal Body Mass Index: A Secondary Analysis of a Randomized Clinical Trial and Observational Study. JAMA Oncol, 2018.

80. Chan, D.S., et al., Body mass index and survival in women with breast cancer-systematic literature review and meta-analysis of 82 follow-up studies. Ann Oncol, 2014. 25(10): p. 1901–14.

81. Iyengar, N.M., et al., Association of Body Fat and Risk of Breast Cancer in Postmenopausal Women With Normal Body Mass Index: A Secondary Analysis of a Randomized Clinical Trial and Observational Study. JAMA Oncol, 2019. 5(2): p. 155–163.

82. Calle, E.E. and R. Kaaks, Overweight, obesity and cancer: epidemiological evidence and proposed mechanisms. Nat Rev Cancer, 2004. 4(8): p. 579–91.

83. Eaton, S., Control of mitochondrial beta-oxidation flux. Prog Lipid Res, 2002. 41(3): p. 197–239.

84. Hoppel, C.L. and S.M. Genuth, Carnitine metabolism in normal-weight and obese human subjects during fasting. Am J Physiol, 1980. 238(5): p. E409–15.

85. Mihalik, S.J., et al., Increased levels of plasma acylcarnitines in obesity and type 2 diabetes and identification of a marker of glucolipotoxicity. Obesity (Silver Spring), 2010. 18(9): p. 1695–700.

86. Markopoulos, C.J., A.K. Tsaroucha, and H.J. Gogas, Effect of aromatase inhibitors on the lipid profile of postmenopausal breast cancer patients. Clinical Lipidology, 2010. 5(2): p. 245–254.

87. Lu, Y., et al., Acetylcarnitine Is a Candidate Diagnostic and Prognostic Biomarker of Hepatocellular Carcinoma. Cancer Res, 2016. 76(10): p. 2912–20.

88. Costell, M., J.E. O’Connor, and S. Grisolia, Age-dependent decrease of carnitine content in muscle of mice and humans. Biochem Biophys Res Commun, 1989. 161(3): p. 1135–43.

89. Kang, M., et al., Metabolomics identifies increases in the acylcarnitine profiles in the plasma of overweight subjects in response to mild weight loss: a randomized, controlled design study. Lipids Health Dis, 2018. 17(1): p. 237.

90. Hansen, J.B., et al., Peroxisome proliferator-activated receptor delta (PPARdelta)-mediated regulation of preadipocyte proliferation and gene expression is dependent on cAMP signaling. J Biol Chem, 2001. 276(5): p. 3175–82.

91. van Vlies, N., et al., PPAR alpha-activation results in enhanced carnitine biosynthesis and OCTN2-mediated hepatic carnitine accumulation. Biochim Biophys Acta, 2007. 1767(9): p. 1134–42.

92. Sen, B., et al., Mechanical strain inhibits adipogenesis in mesenchymal stem cells by stimulating a durable beta-catenin signal. Endocrinology, 2008. 149(12): p. 6065–75.

93. Luu, Y.K., et al., Development of diet-induced fatty liver disease in the aging mouse is suppressed by brief daily exposure to low-magnitude mechanical signals. Int J Obes (Lond), 2010. 34(2): p. 401–5.

94. Krishnamoorthy, D., et al., Marrow adipogenesis and bone loss that parallels estrogen deficiency is slowed by low-intensity mechanical signals. Osteoporos Int, 2016. 27(2): p. 747–56.

95. Luu, Y.K., et al., Mechanical Signals As a Non-Invasive Means to Influence Mesenchymal Stem Cell Fate, Promoting Bone and Suppressing the Fat Phenotype. Bonekey Osteovision, 2009. 6(4): p. 132–149.

96. Yamada, T., et al., Impaired myofibrillar function in the soleus muscle of mice with collagen-induced arthritis. Arthritis Rheum, 2009. 60(11): p. 3280–9.

97. Scheller, E.L., et al., Use of osmium tetroxide staining with microcomputerized tomography to visualize and quantify bone marrow adipose tissue in vivo. Methods Enzymol, 2014. 537: p. 123–39.

98. An, J., et al., Hepatic expression of malonyl-CoA decarboxylase reverses muscle, liver and whole-animal insulin resistance. Nat Med, 2004. 10(3): p. 268–74.

99. Wu, J.Y., et al., ENU mutagenesis identifies mice with mitochondrial branched-chain aminotransferase deficiency resembling human maple syrup urine disease. J Clin Invest, 2004. 113(3): p. 434–40.

100. Jensen, M.V., et al., Compensatory responses to pyruvate carboxylase suppression in islet beta-cells. Preservation of glucose-stimulated insulin secretion. J Biol Chem, 2006. 281(31): p. 22342–51.

101. Xia, J., et al., MetaboAnalyst: a web server for metabolomic data analysis and interpretation. Nucleic Acids Res, 2009. 37(Web Server issue): p. W652–60.

102. Xia, J. and D.S. Wishart, Web-based inference of biological patterns, functions and pathways from metabolomic data using MetaboAnalyst. Nat Protoc, 2011. 6(6): p. 743–60.

103. Xia, J., et al., MetaboAnalyst 2.0--a comprehensive server for metabolomic data analysis. Nucleic Acids Res, 2012. 40(Web Server issue): p. W127–33.

104. Tusher, V.G., R. Tibshirani, and G. Chu, Significance analysis of microarrays applied to the ionizing radiation response. Proc Natl Acad Sci U S A, 2001. 98(9): p. 5116–21.

