## Supplementary figures and images for "Low-Magnitude Mechanical Signals Combined with Zoledronic Acid Reduce Musculoskeletal Weakness and Adiposity in Estrogen-Deprived Mice"

### Supplemental Figure 1

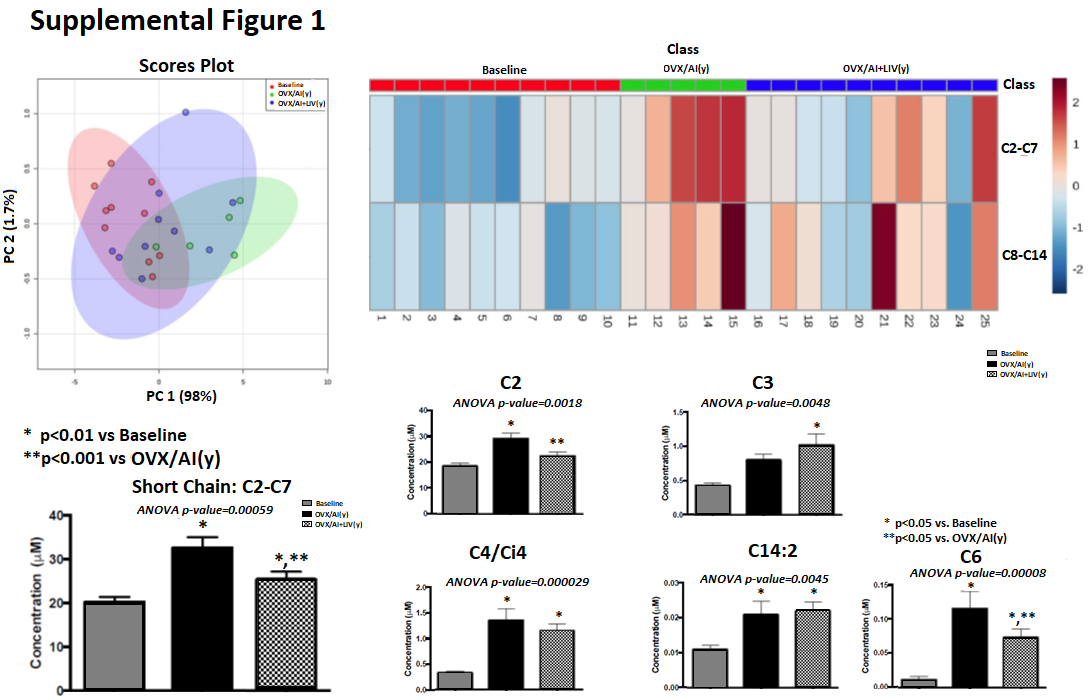

### Supplemental Figure 2

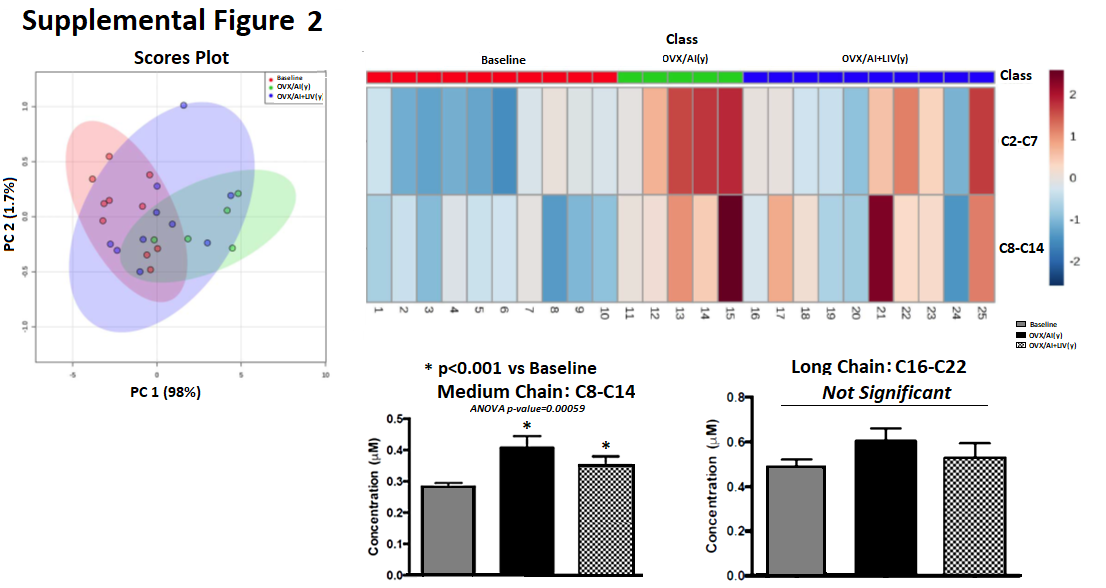

### Supplemental Figure 3

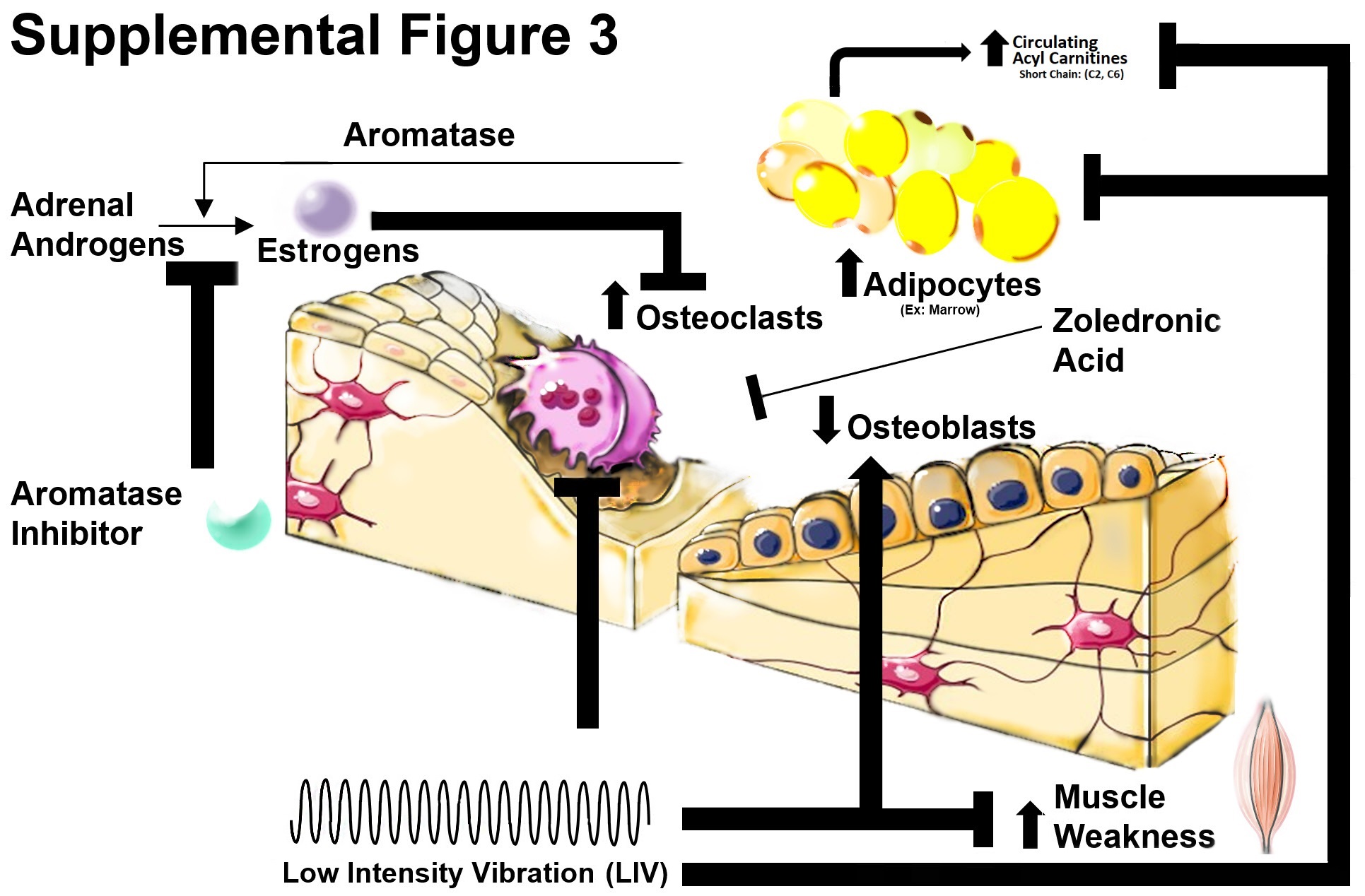

### Supplemental Table 1

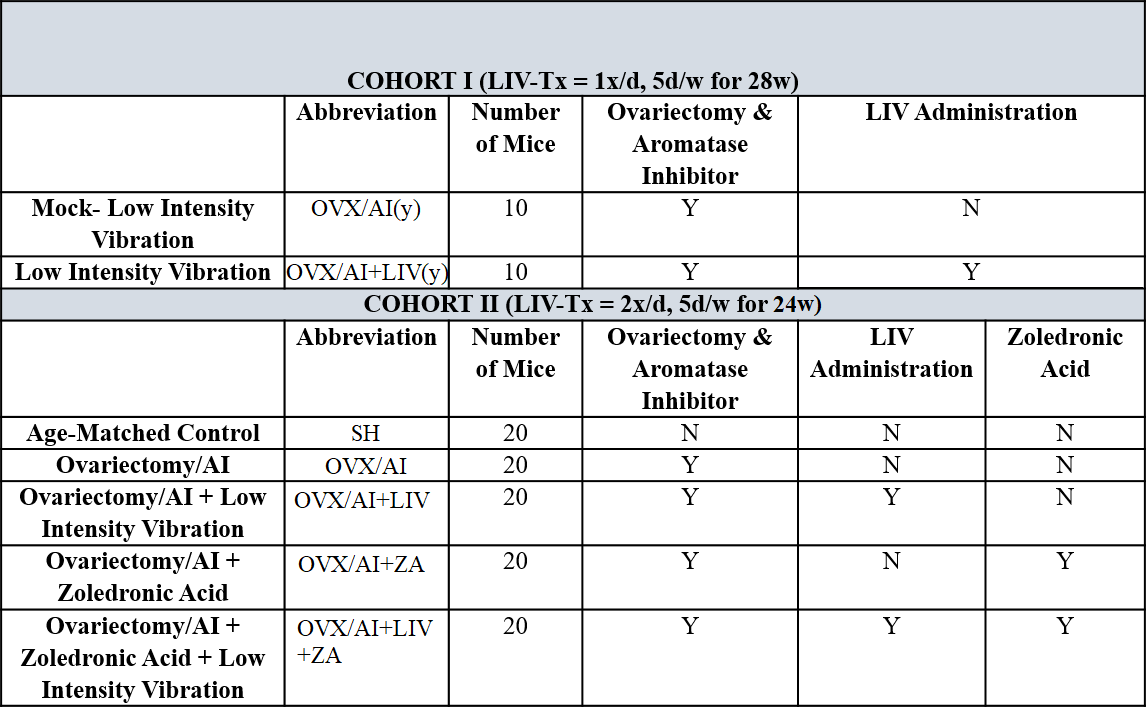

### Supplemental Table 2

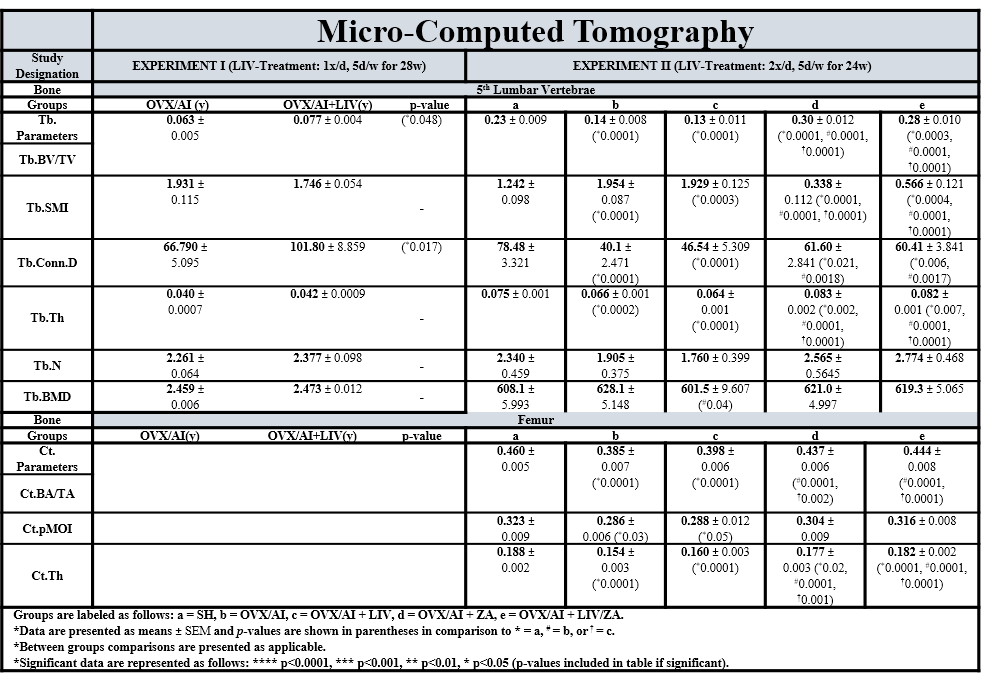

### Supplemental Table 3

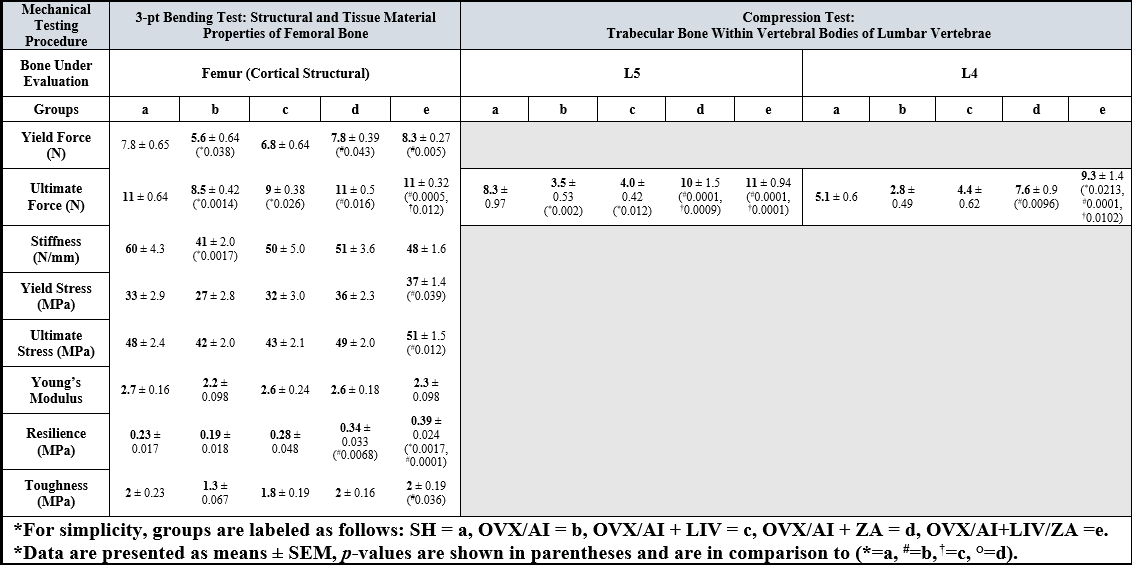
